# Developmental small RNA transcriptomics reveals divergent evolution of the conserved microRNA miR-100 and the *let-7-complex* in nematodes

**DOI:** 10.1101/2024.07.19.604269

**Authors:** Devansh Raj Sharma, Christian Rödelsperger, Waltraud Röseler, Hanh Witte, Michael S. Werner, Ralf J. Sommer

**Affiliations:** Dept. of Integrative Evolutionary Biology, Max Planck Institute for Biology, Tübingen, Germany; School of Biological Sciences, University of Utah, Salt Lake City, Utah, USA

**Keywords:** *Pristionchus pacificus*, *Caenorhabditis elegans*, miRNAs, miR-100, let-7, lin-4, *let-7-complex*, miRNA families, collagens, astacins

## Abstract

**Background:** Over the last three decades, small RNAs have emerged as one of the key post-transcriptional regulators, and are categorised into microRNAs (miRNAs), small interfering RNAs (siRNAs), and PIWI-interacting RNAs (piRNAs). Since their discovery as regulators of developmental timing in the nematode *Caenorhabditis elegans*, some miRNAs have been found to be highly conserved throughout animal evolution. miR-100 is one such highly conserved miRNA, possibly predating the origin of bilaterians. Furthermore, *mir-100* is a member of the conserved *let-7-complex*, a locus containing three conserved miRNA-coding genes (*mir-100*, *let-7*, and *mir-125*/*lin-4*) that does not exist in the same arrangement in *C. elegans*.

**Results:** We performed small RNA sequencing across four developmental time-points in the satellite nematode *Pristionchus pacificus*, identifying 64 novel and ontogenically specifically expressed miRNAs. We also identified miR-100 as the most abundant miRNA in post-embryonic juvenile stages. Additionally, *P. pacificus* exhibited a novel constellation of the *let-7-complex*, wherein *mir-100* is closely linked to *let-7*, but without *mir-125*/*lin-4* in the same locus. Mutants of *mir-100* and *let-7*, individually as well as simultaneously, resulted in viable but developmentally defective worms, showing misregulation in collagen and astacin gene families. Interestingly, *lin-4* mutants showed a delayed but otherwise normal development.

**Conclusion:** This study provides the first developmental small RNA transcriptome of *P. pacificus*, providing a community resource for future evo-devo studies. Furthermore, we identify miR-100 as the most abundant ontogenically-specific miRNA, which, along with let-7, regulates normal development in *P. pacificus*. Lastly, we observe two alternative arrangements of the highly conserved miRNA locus *let-7-complex*.

## Introduction

Development is a complex process, requiring a robust temporal and spatial regulation of gene expression [1–4]. Over the last three decades, small RNAs have emerged as one of the key post-transcriptional regulators of development. Small RNAs can be broadly categorised into microRNAs (miRNAs), small interfering RNAs (siRNAs), and PIWI-interacting RNAs (piRNAs), on the basis of their biogenesis mechanisms [5–8]. These RNAs bind to and guide Argonaute proteins for post-transcriptional repression of target genes [8, 9]. Since their discovery as regulators of developmental timing, first in the free-living nematode *Caenorhabditis elegans* and later in other animals [10–12], small RNAs have been implicated in a wide range of biological pathways, including development and ageing, protection against transposable elements, RNAi, cellular and tissue differentiation, relaying of environmental information, stress response, regulation of access to chromatin, metabolism, antiviral response, and transgenerational inheritance [5, 8, 13–20]. Availability of high-quality genome assemblies and high-throughput sequencing has made studying small RNAs easier in different biological study systems.

Nematodes are arguably the most abundant animal phylum on earth [21] and widely studied in life sciences. While the availability of molecular and genetic tools and resources in *C. elegans* has provided an opportunity to investigate the small RNA molecular machinery in detail [22–31], an increasing number of small RNA studies in other nematodes has widened our understanding of the role of small RNAs in evolutionary and developmental biology, physiology, and in the case of parasitic nematodes, pathology [32–37]. Despite the overarching similarities across the animal kingdom, nematodes show several unique features related to the molecular composition of small RNA-mediated silencing machinery. The most notable anomalies occur in the piRNAs, including their shorter length in comparison to other animals, clustered *vs.* dispersed arrangement in different species, replacement of ping-pong cycle with RdRP-mediated secondary siRNA generation to mediate gene silencing, and the absence of piRNAs altogether in several species [32, 38–43]. Another stark difference is the expansion of the Argonaute protein family along with the apparent loss of DNA methylation machinery in many, but not all, nematodes [44–50]. In this context, the free-living, hermaphroditic diplogastrid nematode *Pristionchus pacificus*, originally developed as a satellite study system for comparative evolutionary developmental studies alongside *C. elegans* [51, 52], is now a model organism in its own right, given the establishment of genetic, genomic, molecular, ecological, behavioural, phylogenetic, morphometric, and biochemical resources [53–60]. Specifically, *P. pacificus* has emerged as a prime model system to investigate developmental plasticity due to its unique feeding polyphenism with two alternative mouth morphs [61–63]. Given that one of the two morphs allows predation on other nematodes, *P. pacificus* is also a new model system to investigate cannibalism, self-recognition, and collective behaviour [64–66]. The repertoire of small RNAs in *P. pacificus* has been examined in a comparative context alongside other nematodes [40–42, 67, 68]. However, the role of small RNAs in *P. pacificus* development has not been investigated.

Here, we report the small RNA transcriptome of *P. pacificus* across multiple developmental timepoints, and identify miR-100 as ontogenically specifically the most abundantly expressed miRNA. Subsequent analyses identified that miR-100, the most ancestrally conserved miRNA in the animal kingdom, is tightly genetically linked to the miRNA let-7 in the *P. pacificus* genome. This genetic linkage is partially reminiscent of the *let-7-complex* [69, 70], a conserved genetic locus of miRNA-coding genes which does not exist in its canonical form in *C. elegans*. Instead, *C. elegans* encodes 6 divergent miRNAs that share the same seed sequence with miR-100. Genetic mutants of the two miRNAs showed pleiotropic phenotypes pertaining to developmental defects and reduction in brood size. RNA sequencing of these mutants showed that the two miRNAs ensure normal development by regulating transcription of collagens and astacins, two large protein classes found in nematodes that regulate various aspects of development, such as fertilisation, hatching, and periodic cuticle building and shedding, among others.

## Methods

### Stage-specific worm pellet collection

10-cm NGM-agar plates spotted with 800 •l *E. coli* OP50 were seeded with *P. pacificus* wild-type strain (PS312) and incubated at 20°C [71]. Fully populated mixed-stage plates were washed down 6 days post seeding using M9, then filtered through a 5 µm nylon filter to separate bacteria (in the flow-through) and worms (on the filter). Filtered worms were collected and bleach-treated to rid the mixture of all post-hatched worms and any remaining bacteria [71], followed by a filtration through a 120 µm nylon filter to separate eggs (in the flow-through) from any undissolved adult worm carcasses (on the filter). The flow-through was gently centrifuged (500g for 2 minutes) to collect the egg pellet and remove the supernatant, followed by washing and resuspension in M9. Approximately 30% of the egg resuspension volume was transferred to microcentrifuge tubes to collect the embryo developmental stage (Fig. 1A) by centrifugation followed by flash-freezing in liquid nitrogen and storage at −80°C. Remaining egg resuspension was spotted on to empty 10-cm NGM-agar plates with OP50 for collection of older developmental stages. Three (including pyrophosphatase treatment) or two (excluding pyrophosphatase treatment) biological replicates were collected per developmental time point.

**Fig. 1:**
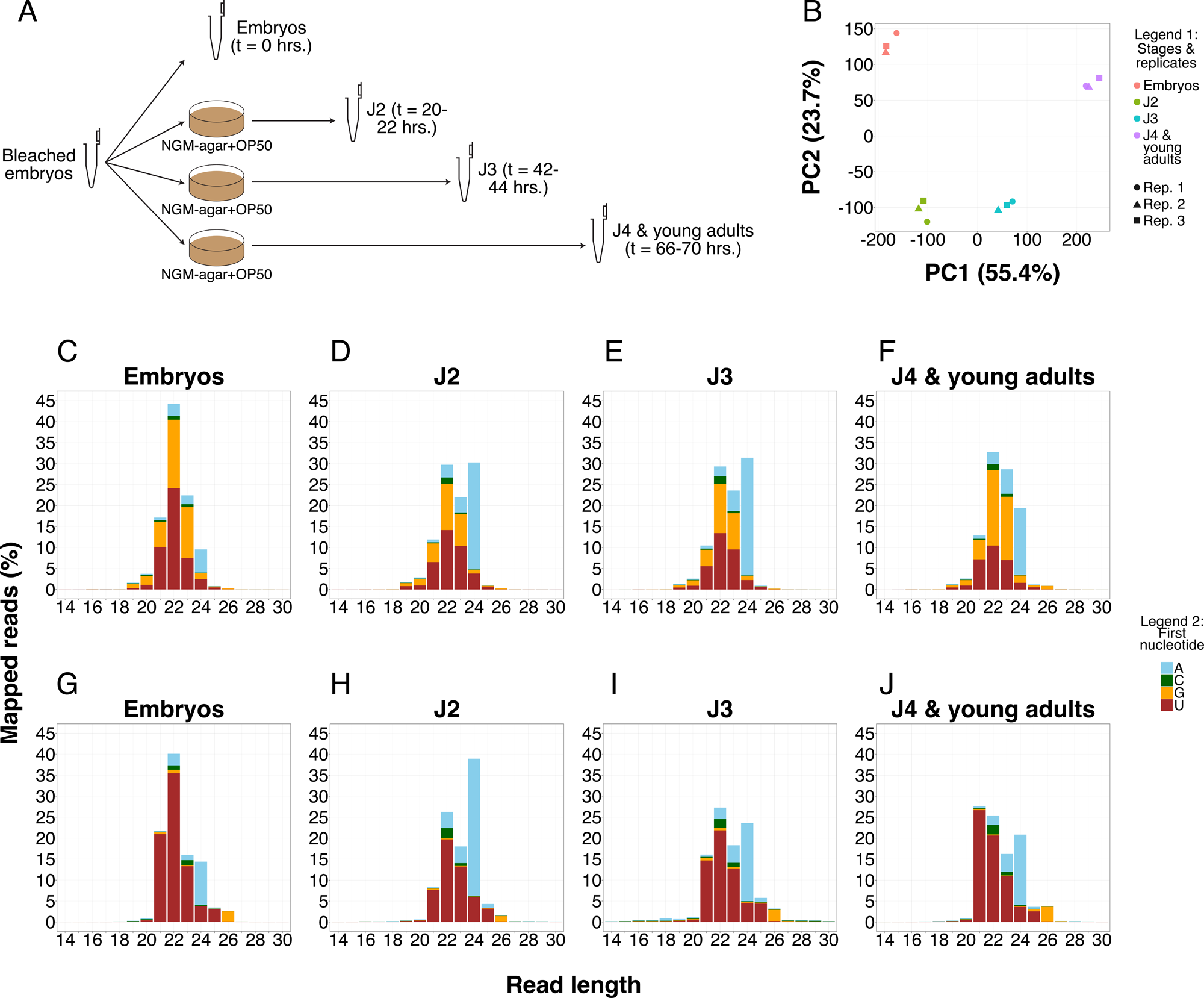
Developmental small RNA profile of *P. pacificus*: A: Schematic representation of experimental design. B: Principal component analysis (PCA) of the developmental stages of the small RNA transcriptome. Colours correspond to different developmental stages, and shapes correspond to different biological replicates per developmental stage (Legend 1). C–F: Small RNA profile at different developmental stages, categorised on the basis of read length (x-axis) and first 5’ nucleotide (Legend 2). Y-axis denotes mean abundance of each category as a fraction of whole small RNA transcriptome for that corresponding stage. G–J: Small RNA profile at different developmental stages without pyrophosphatase treatment.

For J2 pellet collection, plates spotted with bleached eggs were washed off 20–22 hours post seeding, using M9 (Fig. 1A). Washed worms were first filtered through a 20 µm nylon filter to separate J2 and bacteria (in the flow-through) from unhatched and dead embryos (on the filter), followed by filtering the flow-through using a 5 µm nylon filter to separate bacteria (in the flow-through) and worms (on the filter). For J3 pellet collection, bleached eggs were washed off 42–44 hours post seeding (Fig. 1A). Washed worms were filtered through a 20 µm nylon filter to separate J2 and bacteria (in the flow-through) from J3 worms (on the filter). For J4/young adult (YA) pellet collection, bleached eggs were washed off 66–70 hours post seeding (Fig. 1A). Washed worms were filtered through a 5 µm nylon filter to separate J4/YA worms (on the filter) from bacteria (in the flow-through). In all cases, worms were washed off the filter patch and resuspended using M9, transferred and centrifuged in microcentrifuge tubes to collect into a pellet and flash-frozen in liquid nitrogen before being stored at −80°C.

For analysis of wild-type *vs.* mutants, J3 pellets were collected post bleaching as described above. Three (for *Ppa-mir-100*) or two (for the rest) biological replicates were collected per strain.

### Total RNA extraction, small RNA size selection, NGS library preparation, and sequencing

Worm pellets as collected above were thawed on ice and added with 500 µl of TRIzol (Invitrogen), followed by a quick vortex treatment for resuspension of pellets. Next, three cycles of flash-freezing the worm-TRIzol mix in liquid nitrogen, followed by thawing at 37°C, were performed. Following three freeze-thaw cycles, 100 µl of chloroform was added to the worm-TRIzol mix, and the contents of tubes mixed by shaking and inverting. Microcentrifuge tubes containing the worm-TRIzol-chloroform mix were incubated at room temperature for 2–3 minutes, followed by a centrifugation at 12000g and 4°C for 15 minutes. Post centrifugation, the aqueous phase on top was transferred to a new sterile microcentrifuge tube for each sample, equal volume (approximately 300 µl) of 100% ethanol was added, and the contents of tubes mixed by vortexing. Total RNA from the aqueous phase-ethanol mix was subsequently extracted using RNA Clean & Concentrator-25 kit (Zymo Research) following the manufacturer’s protocol. Extracted RNA was then subjected to CapClip acid pyrophosphatase (CCAP) (Biozym) treatment for removal of 5’ triphosphate cap (exception: 5’ monophosphate-biased library, where this step was skipped; Fig. 1G–J). For pyrophosphatase treatment, total RNA was sequentially added with sterile-grade water, 5 µl of 10x reaction buffer (Biozym), and lastly with 5 µl of CCAP enzyme (Biozym) in a microcentrifuge tube over ice (total reaction volume = 50 µl). Contents of the tubes were then mixed by gentle pipetting, followed by incubation at 37°C for 2 hours, followed by a clean-up step. RNA concentration was determined using the Qubit fluorometer (Invitrogen) before and after pyrophosphatase treatment for assessment of RNA losses.

Following clean-up, RNA was used for preparation of small RNA next generation sequencing libraries with NEBNext Multiplex Small RNA Library Prep Kit (E7560S, New England BioLabs Inc.) using the manufacturer’s protocol, including a negative (no RNA) control. RNA libraries thus prepared were subjected to size selection using Blue Pippin and compliant 3% agarose gel cassettes (Sage Science Inc.) following the manufacturer’s protocol. RNA samples before library preparation, and NGS libraries after size selection and a DNA cleanup step (using Monarch PCR & DNA cleanup kit; T1030S, New England BioLabs Inc.) were analysed with Bioanalyzer (Agilent Technologies, Inc.) using the manufacturer’s protocol to check for integrity of RNA, library size, and quality. Once satisfactory profiles were observed on Bioanalyzer, libraries were pooled and run on a single lane on an Illumina HiSeq 3000 machine.

For J3 stage-specific small RNA sequencing of wild-type and mutants, total RNA was extracted as described above. One part of total RNA was kept for RNA sequencing, while the other part was treated with pyrophosphatase enzyme for small RNA sequencing. Both types of samples were then submitted to Novogene (UK) Company Ltd. for RNA and small RNA sequencing respectively.

### Analysis of small RNA-seq data

Small RNA sequencing raw reads were first subjected to adapter trimming using Cutadapt (version 4.1) [72], then filtered for reads mapping to *P. pacificus* tRNA and rRNA genes identified through tRNAscan-SE (version 2.0.5) and RNAmmer (version 1.2) respectively [73–75]. Remaining reads were then mapped to the *P. pacificus* genome using Bowtie2 (version 2.3.5.1) [57, 76, 77], procuring the FASTA sequences from BAM alignments using SAMtools (version 1.10) [78, 79]. FastQC (Babraham Bioinformatics) was used for ensuring quality of sequences and presence or absence of adapters.

Custom shell scripts were written for sorting reads on the basis of length and first 5’ nucleotide, the output of which was fed to R (versions 4.2.3 and 4.5.2) to generate small RNA profiles (Fig. 1C–J, Fig. 5A–D). For identifying the putative Ruby motif in *P. pacificus*, using existing knowledge about *P. pacificus* piRNAs [40], we queried for top 5 motifs 70– 100 base-pairs upstream of 21U reads using MEME [80, 81] from the MEME suite (version 5.2.0). For all predicted motifs for each of the 4 developmental timepoints, the best motifs predicted had a much better E-value (E-value < 10^-4^) than the second best (E-value ∼ 0.1) and closely resembled the Ruby motif. Using the best predicted motifs for each developmental stage as query, their occurrence was searched for 100 base-pairs upstream of all small RNA reads between lengths 19 and 26 nucleotides using FIMO [82] from the MEME Suite, followed by looking at the length (Fig. S1A–D) and first 5’ nucleotide of positive sequences. For miRNAs, we looked for sequences with perfect matches to the mature miRNAs FASTA sequence from miRBase [83]. Remaining sequences were categorised as ‘others’ (Fig. S1A–D).

Principal component analysis was done with individual biological replicates of the developmental small RNA transcriptome (Fig. 1B) and the J3 stage-specific small RNA sequencing of wild-type and mutants (Fig. S5C) using the ‘prcomp()’ function in R. Read counts were normalised per million, and sequences with an average below 1 CPM were filtered out of the analysis.

Differential expression analysis was done for small RNAs between the four developmental stages, as well as between the wild-type and *Ppa-mir-100* alleles, in a pairwise manner in R using DESeq2 (version 1.50.2) [84], using small RNA sequences with a mean count • 10 across the two developmental stages/genotypes per pairwise comparison. We always included a log_2_-fold change cut-off of 1 (absolute value; corresponding to a fold-change of 2, or 100% change in expression levels) into our null hypothesis. Differentially expressed small RNAs (DEsRNAs) with an adjusted p-value < 0.1 were short-listed for inclusion in our results (Tables S31–S36, S38–S43).

### Nomenclature of microRNAs

Gene names for miRNA-coding genes in *P. pacificus* carry the species-corresponding prefix ‘*Ppa-*’, similar to the gene nomenclature of protein-coding genes and in accordance with the 3-letter species-specific abbreviation designated to *P. pacificus* within the nematode community. In contrast, the corresponding miRNAs carry the species-corresponding prefix ‘ppc-’, in accordance with the 3-letter species-specific abbreviation designated to *P. pacificus* on miRBase. Furthermore, miRNAs are denoted in non-italicised text, and with an uppercase ‘R’ if the name contains the prefix ‘miR-’, while their corresponding miRNA-coding genes are italicised and with a lowercase ‘r’ in the prefix ‘*mir-*’ in accordance with the miRNA nomenclature conventions [85, 86]. Thus, ‘ppc-miR-100’, ‘ppc-lin-4’, ‘ppc-let-7’, and ‘ppc-miR-2235a-3p’ are the miRNAs corresponding to the miRNA-coding genes ‘*Ppa-mir-100*’, ‘*Ppa-lin-4*’, ‘*Ppa-let-7*’, and ‘*Ppa-mir-2235a-3p*’ respectively.

### Identification of novel miRNAs with miRDeep2

For identification of novel miRNAs across the four developmental stages, small RNA sequencing raw reads generated in this study as well as those generated for *P. pacificus* in a previous study [41] (since we suspected that novel miRNAs from this particular study were not present on miRBase; please also see ‘Results and discussion’) were subjected to analysis through miRDeep2 [87, 88]. As miRDeep2 includes a module for predicting novel miRNAs for the species in question, as well as provide a metric, called ’miRDeep2 score’, as an indicator of the support for those predictions, we analysed the output .csv files generated by miRDeep2 analysis of the sequencing reads pertaining to the four developmental stages from this study, as well as the four conditions (TAP-treated adults, TAP-treated dauers, untreated adults, and untreated dauers) from the previous study [41], in R to ascertain novel miRNAs in both studies. Using a cut-off of 2 for miRDeep2 score, and removing those sequences that were also identified as novel miRNAs in the previous study [41] from our datasets, we identified novel miRNAs in this study (Fig. S1E, Table S4). Developmental stage-specific abundance of the novel miRNAs was then identified by calculating the proportion of mature read count — as determined by miRDeep2 — for those miRNAs in the total number of small RNA transcriptome reads for the corresponding developmental stages, averaged over three biological replicates (Fig. S1F– I).

For identification of the most abundant reads, for each category (combination of read length and first 5’ nucleotide) of small RNAs, the number of reads pertaining to each unique genome-mapped small RNA sequence were counted between lengths 19 and 26 using custom Shell scripts. The resulting tables were fed into R to identify reads accounting for 1% or more of the corresponding category (Fig. 2A–C, Fig. S2A).

**Fig. 2:**
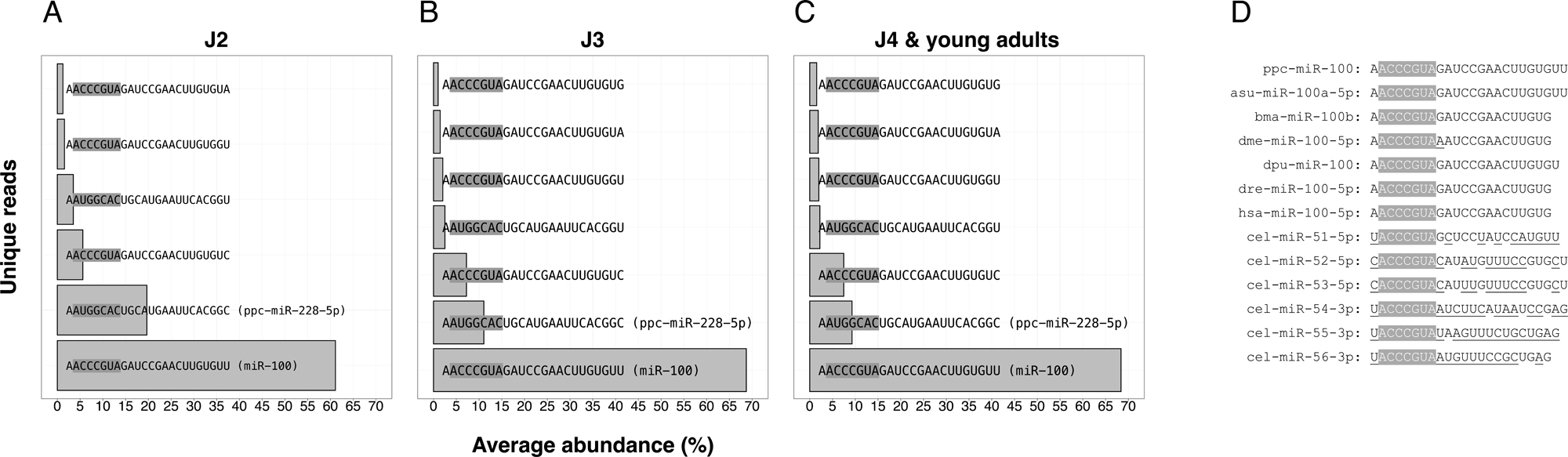
Abundance of ppc-miR-100, a highly conserved miRNA across animal kingdom, in juvenile developmental stages of *P. pacificus*: A–C: Most abundant 24A reads (> 1%) in three developmental stages, represented as a fraction of total 24A reads, arranged in ascending order of abundance from top on y-axis. Read#1, with sequence AACCCGUAGAUCCGAACUUGUGUU, corresponds to a highly conserved metazoan miRNA known as miR-100 (seed sequences are highlighted; note that the lesser abundant sequences appear to be variants of miR-100 and ppc-miR-228-5p, identified by their common seed sequence and 5’ end but differing at their 3’ end, usually in a SNP. As these sequences do not have a perfect match in the *P. pacificus* genome, we cannot ascertain if these sequences represent sequencing errors or post-transcriptional modifications). D: Comparison of miR-100 sequences from different species. Note the sequence divergence outside of the seed region in *C. elegans* in comparison to other species (seed sequence is highlighted, and sequence alterations are underlined) (species abbreviations: ppc; *P. pacificus*, asu; *A. suum*, bma; *B. malayi*, dme; *D. melanogaster*, dpu; *D. pulex*, dre; *D. rerio*, hsa; *H. sapiens*, cel; *C. elegans*)

### Quantification of expression levels

For the expression levels of the three miRNAs miR-100, let-7, and lin-4 in *P. pacificus*, the number of reads with perfect match to the three sequences (ppc-miR-100: AACCCGTAGATCCGAACTTGTGTT, ppc-let-7: TGAGGTAGTAGGTTGTATAGTT, ppc-lin-4: TCCCTGAGACCTCAATTGTG) were identified, across the corresponding developmental stages or mutant strains (Fig. 3D–F, Fig. 5M–N, Fig. S5A–B).

**Fig. 3:**
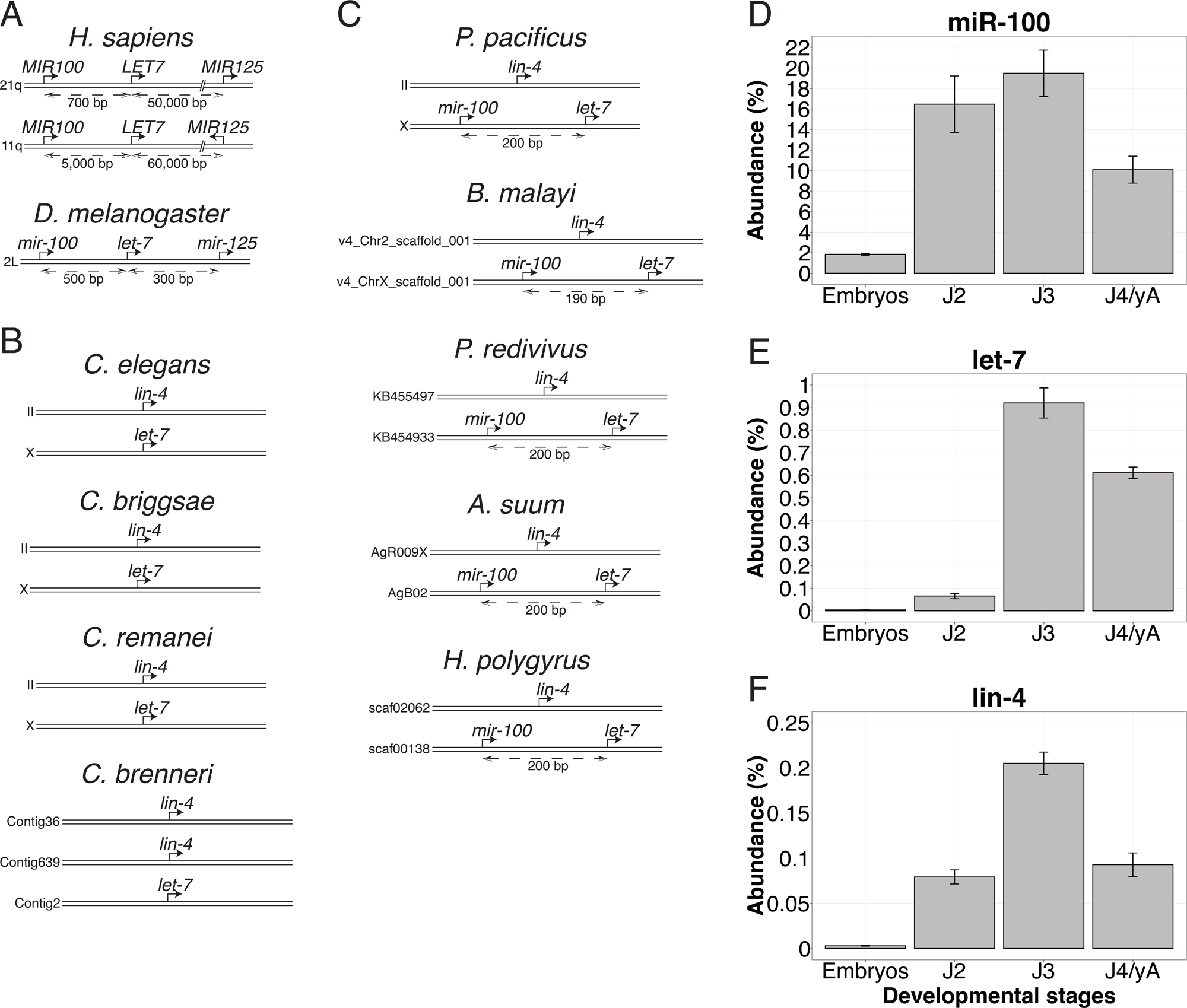
Two alternative arrangements of *let-7-complex* in nematodes: A: In humans, fruit flies, and other animals, miRNA-coding genes *mir-100*, *let-7* and *mir-125*/*lin-4* are found on the same chromosome in a genomic locus called the *let-7-complex*. Note that the miRNA-coding genes are denoted using the corresponding species-specific miRNA nomenclature conventions as mentioned in [86]. B: *Caenorhabditis* nematodes show a complete dissociation of the *let-7-complex*, with *lin-4* and *let-7* on two different chromosomes, along with an expansion and divergence of ancestral *mir-100* into six miRNA-coding genes (see Fig. 2D and main text). C: *P. pacificus* and other non-*Caenorhabditis* nematodes investigated in this study show an alternative arrangement, in which ancestral *mir-100* is retained in the genome, and is found upstream of *let-7* on the same chromosome in close proximity, but *lin-4* is found on a different chromosome. D–F: Developmental expression profiles of miRNAs miR-100, let-7 and lin-4 in *P. pacificus*. All three miRNAs are at their developmental peak in J3. Bar plots represent the mean abundance of the corresponding miRNA as a percentage of the entire small RNA transcriptome, and error bars represent the SEM, over three replicates. Note that the y-axis is different for the 3 miRNAs.

For miRNA family expression levels, custom Shell scripts were written to identify the number of sequences matching the seed sequence of corresponding miRNAs (nucleotides 2–8 [89]) across the mutant strains (Fig. 5E–L).

### Analysis of RNA-seq data

Paired-end RNA sequencing reads were aligned against the *P. pacificus* genome using STAR (version 2.7.9a) [57, 90], and mapped sequences were counted against the *P. pacificus* gene annotations (El Paco gene annotations, version 3) using featureCounts from the Subread package (version 2.0.0) [91, 92]. Differential expression analysis was done between wild-type and mutant strains in a pairwise manner in R using DESeq2 (version 1.38.3) [84], using genes with a mean count • 10 in the wild-type strain. We always included a log_2_-fold change cut-off of 0.5849625 (absolute value; corresponding to a fold-change of 1.5, or 50% change in expression levels) into our null hypothesis. Differentially expressed genes (DEGs) with an adjusted p-value < 0.1 were selected for subsequent domain enrichment analyses. The latter were based on two-tailed Fisher’s Exact tests with FDR-correction of p-values. To explore the temporal expression pattern of the DEGs, we visualised their expression level using a previously generated high-resolution transcriptomic dataset [93].

DEGs were further analysed to gain insights on their biology by being subjected to protein domain enrichment analysis and overrepresentation analysis, the latter in context of two published genome-wide datasets categorising *P. pacificus* genes into co-expression modules and their oscillating nature [93, 94]. For protein domain enrichment analysis, we identified the enrichment and corresponding Rich Factor for protein domains in DEGs (Fig. 6E–J). Enrichment of a domain X is a ratio of ratios, defined as: (DEGs with domain X ÷ DEGs without domain X) ÷ (non-DEGs with domain X ÷ non-DEGs without domain X); Rich Factor of a domain X is a measure of its degree of enrichment, and is defined as: (DEGs with domain X ÷ all genes with domain X). For overrepresentation analysis, we compared if our DEGs were overrepresented in comparison to the reference distribution of genes across co-expression modules [94] and in comparison to the percentage of oscillating genes [93] across the *P. pacificus* genome. P-values were calculated using Fisher’s Exact test with FDR correction for multiple testing.

**Fig. 4:**
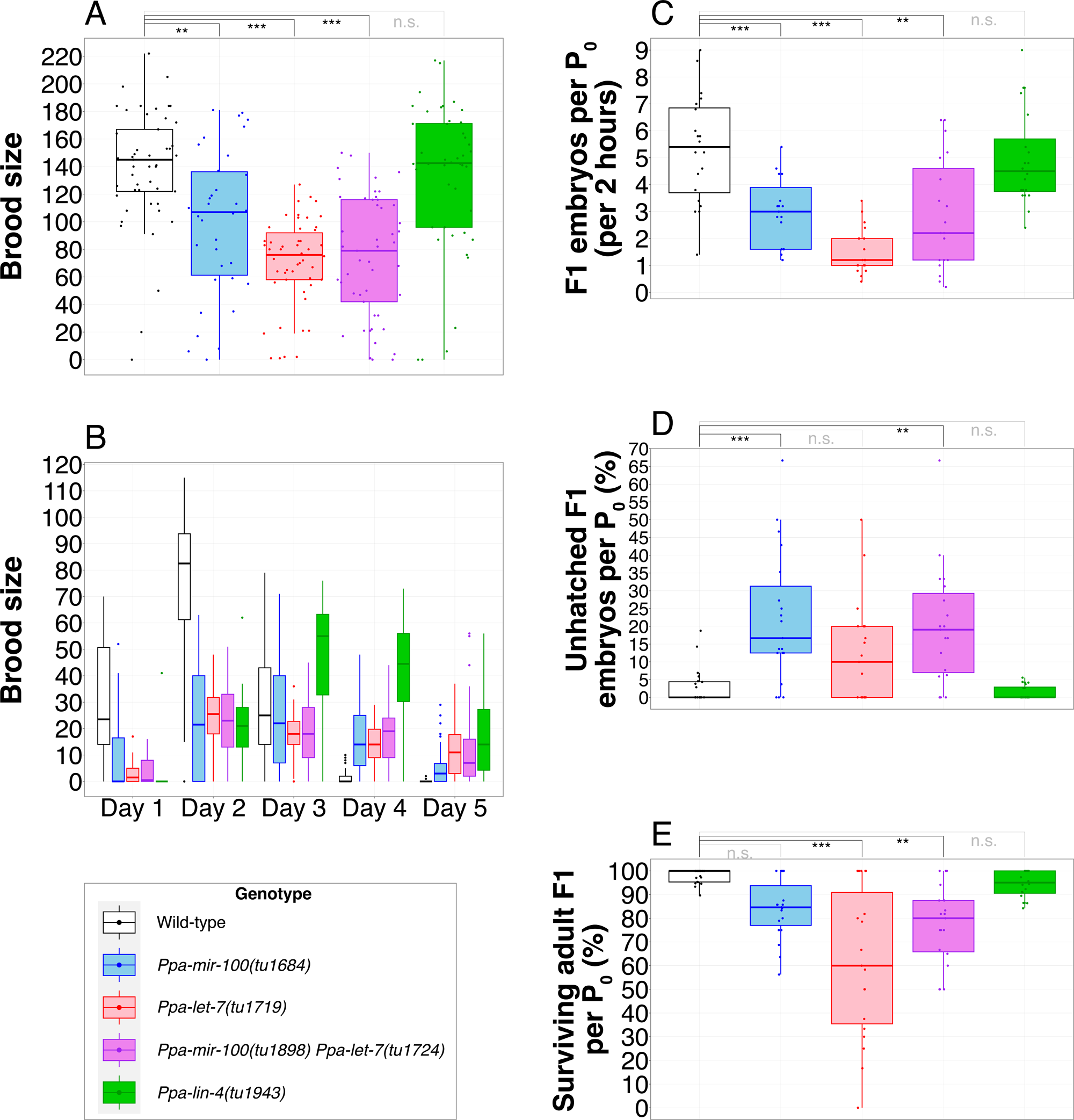
Phenotypic characterisation of *Ppa-mir-100*, *Ppa-let-7*, *Ppa-mir-100 Ppa-let-7* double mutant, and *Ppa-lin-4* mutants: A–B: Brood sizes of different genotypes, taken all together (A) or on a temporal basis (B). n = 50 mothers per genotype for A and B. C: Frequency of egg-laying over 2 hours per synchronised mother for each genotype. D: Embryonic lethality per synchronised mother per genotype. E: Percentage of surviving embryos that reach adulthood per synchronised mother per genotype. n = 100 mothers per genotype for panels C–E. Statistical significance of mutant phenotypes against wild-type strain for panels A, C, D, and E are shown above the corresponding panels. For statistical significance of all pairwise comparisons in all panels, see Table S8

**Fig. 5:**
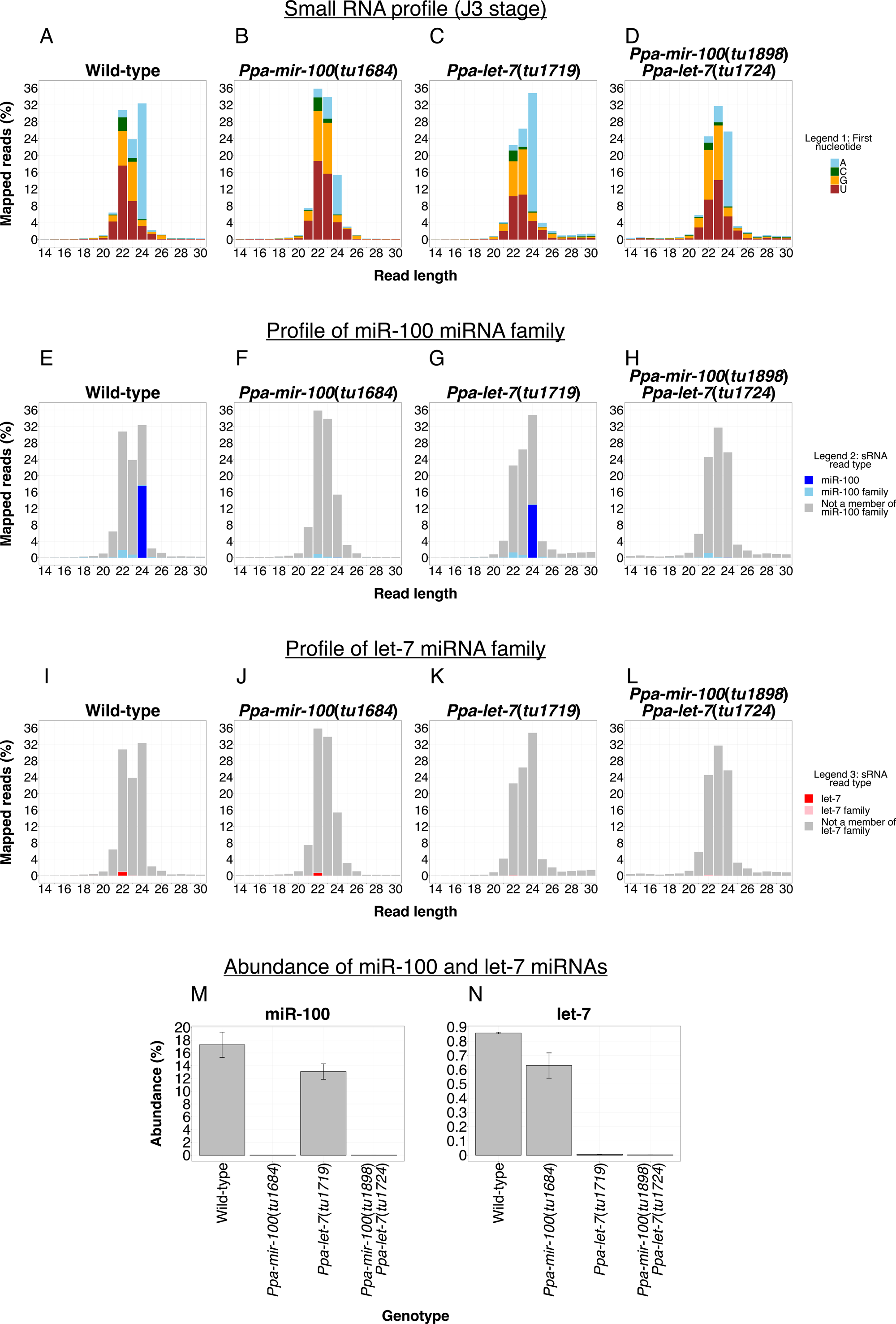
Small RNA sequencing of wild-type, *Ppa-mir-100*, *Ppa-let-7* and *Ppa-mir-100 Ppa-let-7* double mutants of *P. pacificus*: A–D: Small RNA profile of the four genotypes at J3 stage, categorised on the basis of their length (x-axis) and first 5’ nucleotide (Legend 1). Y-axis denotes the mean abundance of each category as a fraction of the whole small RNA transcriptome for that corresponding genotype. E–L: Profile of miR-100 family of miRNAs (E–H) or of let-7 family of miRNAs (I–L) for the corresponding genotypes as in panels in A–D. Legend 2 corresponds to panels E–H, Legend 3 to panels I–L. M–N: Expression levels of miRNAs miR-100 (M) and let-7 (N) across the four genotypes. Error bars represent the SEM. Note that the y-axis is different between panels M and N.

**Fig. 6:**
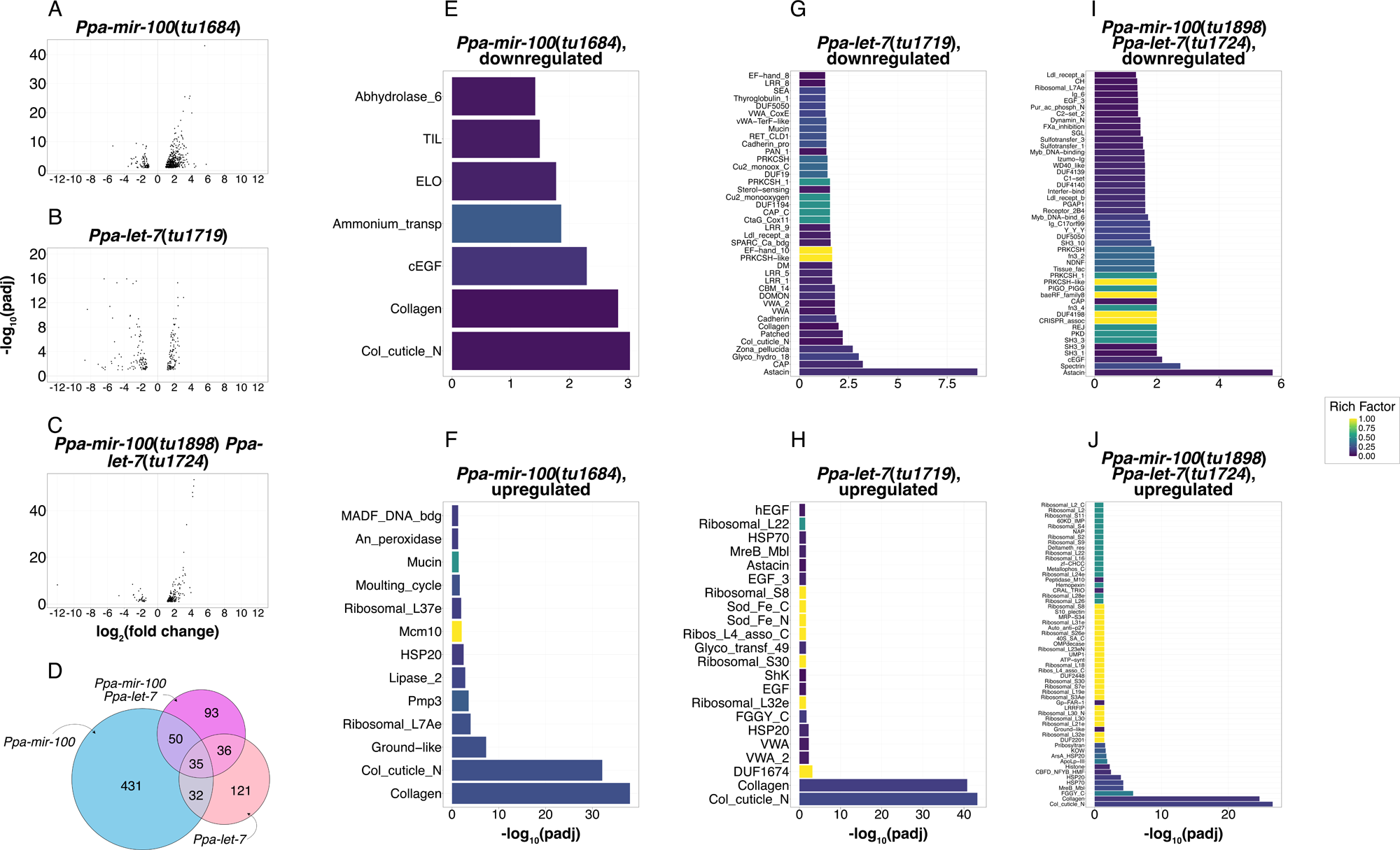
RNA sequencing of wild-type, *Ppa-mir-100*, *Ppa-let-7*, and *Ppa-mir-100 Ppa-let-7* double mutants of *P. pacificus*: A–C: Volcano plots of the three mutant genotypes at J3 stage, resolved along fold-change compared to wild-type (x-axis) and adjusted p-value (y-axis). Note that the y-axis is different between panels A–C. D: Venn diagram showing the number of overlapping genes between the three mutants. E–J: Domain enrichment analysis of differentially expressed genes in the three mutant genotypes, separated into two plots for each genotype for downregulated (E, G, I) and upregulated (F, H, J) genes, showing all statistically significantly enriched (padj < 0.05) domains on y-axis, their corresponding adjusted p-values on x-axis, and their Rich Factor represented by the colour scale. Note that the x-axis is different between panels E–J.

### Identification of putative miRNA targets

To identify putative direct mRNA targets of miRNAs, we first defined the region 100 base-pairs upstream till 200 base-pairs downstream of the end of an annotated *P. pacificus* protein (El Paco gene annotation, version 3) [92] as presumptive 3’ UTRs corresponding to those genes. We then queried for perfect complements (5’-ACGGGT-3’ for miR-100, 5’-TACCTC-3’ for let-7) to the seed region of the miRNAs, taking into account nucleotides 2– 7 instead of 2–8 to procure a more inclusive list of putative mRNA targets (Tables S6–S7).

### Data visualisation

Data wrangling for graphical representation was done in R using ggplot2 (version 3.4.3) [95]. The Venn diagram (Fig. 6D) was prepared using the eulerr package (version 7.0.0) [96]. Post plotting, aesthetic modifications were performed in Adobe Illustrator CS6 (version 16.0.4) and Affinity Designer 2 (version 2.6.5).

### Phylogenetic analysis

To investigate the evolutionary history of the miR-100 family in nematodes, we employed the search function of miRBase website to screen for other members of miR-100 family across various nematodes. The resulting hits were aligned using MUSCLE (version 5.2) [97], yielding a multiple sequence alignment of 24 nucleotide positions. A maximum-likelihood tree was calculated by the phangorn R package using the F81+I model, which was selected as the best model based on the Bayesian information criteria [98]. In addition, we calculated all pairwise differences from the multiple sequence alignment and visualised the results as a heat-map. Sequences were arbitrarily rearranged and grouped after visual inspection of the patterns of pairwise differences.

### CRISPR/Cas9-mediated mutagenesis

CRISPR/Cas9-mediated mutagenesis was performed based on previously published protocols with some modifications [56, 58, 99, 100]. In short: crRNAs were designed with high sequence specificity 20 base-pairs upstream of the protospacer adjacent motif (PAM) sites (Integrated DNA technologies, Inc.). Cas9 protein (Catalog no. 1081058), tracrRNA (Catalog no. 1072534) and all three crRNAs were purchased from Integrated DNA Technologies. 5 µl of tracrRNA (0.4 µg/µl) and 1.12 µl of crRNA (0.4 µg/µl) were mixed and incubated at 37°C for 10 minutes followed by a 5-minute incubation at room temperature. To the above mix, 0.5 µl of Cas9 protein (10 µg/µl), 1 µl (100 mM) of 100-nucleotide long DNA repair template carrying a deletion for the miRNA sequence and 3 µl (170 ng/µl) of plasmid PZH009 [58] carrying the promoter of *Ppa-eft-3* coupled with TurboRFP [101] as co-injection marker were added, and the total reaction volume made up to 20 µl using Tris-EDTA buffer. Injected worms (P_0_) were put onto NGM-plates with 50 µl OP50 lawns and allowed to lay eggs at 20°C for 24–32 hours before being sacrificed. Three to four days later, F1 J4s/YAs were screened for RFP co-injection marker fluorescence, and those containing RFP-expressing F1s were segregated onto individual plates, and genotyped after ∼24 hours. F2 progeny of heterozygous F1s were genotyped (typically 10– 16 F2s per F1) for potential homozygous mutants. Upon identification of such homozygous F2s, their progeny (F3) were subjected to the above procedure for a confirmative Sanger sequencing (typically 4–6 F3s per homozygous F2), following which mutant lines were established. Sanger sequencing was done by Azenta Life Sciences. Allele names and molecular lesions of all mutants generated in this study are mentioned in Fig. S7.

### Phenotypic assays

A schematic layout is provided in Fig. S4. Briefly, day 1 adult hermaphrodites, with no more than 1–2 eggs in the uterus, from non-starved stock maintenance plates for 5+ generations were seeded to empty NGM-agar plates with 300 µl OP50 lawns, and allowed to lay eggs for 2 hours, after which they were removed, leaving the plates with 2-hour synchronised P_0_ worms (maternal synchronisation; Fig. S4). For the egg-laying and survival assay, these synchronised P_0_ young adult worms, after 90–96 hours, were passed onto NGM-agar plates with fresh (<24 hours old) 50 µl OP50 lawns (5 worms per plate) and allowed to lay eggs for 2 hours at room temperature before being removed. The number of synchronised F1 eggs were scored for egg-laying rate (Fig. 4C, Fig. S3A–C), before being incubated at 20°C. Approximately 28–30 hours later, the incubated F1 embryos were scored for embryos still unhatched (Fig. 4D), before being incubated again at 20°C. 90–96 hours post F1 synchronisation, the number of surviving F1 adults were scored (Fig. 4E). A total of five biological replicates (20 P_0_ worms per replicate) were performed per strain.

For brood size assay, starting from 2-hour synchronised P_0_ embryos, these embryos were incubated at 20°C for ∼70 hours, by which time they would have developed into the J4 stage. At this stage, P_0_ worms were singly segregated onto individual “day 1” plates and incubated at 20°C. After 24 hours, these worms were seeded to new “day 2” plates, and both sets of plates were incubated at 20°C. The experiment continued till accumulation of “day 5” plates. On day 6, P_0_ worms on “day 5” plates were sacrificed and the plates incubated at 20°C, and the “day 1” plates were scored for the number of F1 adult worms (Fig. 4B). 24 hours later (day 7), “day 2” plates were scored, and so on until day 10, when “day 5” plates were scored for F1 adults. Total brood size was calculated by summing up the per day scores (Fig. 4A). Two replicates (25 P_0_ worms per replicate) were performed per strain.

To test for mutant phenotypes in our assay data, we applied a permutation-based version of MANOVA in which null model residuals were randomised (in 10,000 iterations) using the RRPP package (version 1.4.0) [102]. P-values for the pairwise comparisons of individual strains are reported in the corresponding figure panels (Fig. 4, Fig. S3) and in tabular form (Table S8) (*: p-adj < 0.05, **: p-adj < 0.01, ***: p-adj < 0.001). We considered our data as incompatible with the null hypothesis (i.e., “statistically significant”) if the derived (FDR-corrected) *P*-values were < 0.05.

## Results and Discussion

### Stage-specific small RNA sequencing reveals developmental dynamics in *P. pacificus*

To investigate the small RNA transcriptome of *P. pacificus*, worms from four synchronised time points were collected, covering five developmental stages: embryos, J2s, J3s, and late J4s with young adults (YA) (3 biological replicates per time point) (Fig. 1A). Nearly all small RNAs were between 19 and 26 nucleotides in length, and began with either a uracil, guanine, or adenine (Fig. 1C–F). This observation is in line with the previous analyses of *P. pacificus* small RNAs, and with studies showing a 5’ bias towards adenine or uracil in miRNAs and piRNAs, and towards guanine in primary and secondary siRNAs in *C. elegans* [23, 25, 26, 41, 103–107]. Additionally, a second set of sequencing experiments was performed without pyrophosphatase treatment (2 biological replicates per time point) (Fig. 1G–J), to exclude 5’ triphosphate and capped-small RNAs, in particular, RdRP-generated secondary siRNAs, while retaining 5’-monophosphate bearing miRNAs, piRNAs, and primary siRNAs [23, 40, 41, 104, 105]. Comparing the corresponding stages between the two experiments (Fig. 1C–J) showed depletion in small RNAs with a 5’ guanine, except for reads with length of 26 nucleotides, corresponding to primary siRNAs. This finding is in accordance with previous knowledge of secondary siRNAs having a bias towards 5’ guanine and length 22–23 nucleotides [26, 41, 104]. Lastly, we also performed a principal component analysis (PCA) with our pyrophosphatase-treated small RNA datasets to confirm reproducibility of data between our biological replicates. PCA showed that the 12 samples clustered among themselves on the basis of their corresponding developmental stages, along with a clear segregation between the different clusters obtained (Fig. 1B). Thus, the PCA confirmed reproducibility among replicates of our generated data, in addition to confirming that the distinction between small RNA profiles at different developmental stages (Fig. 1C–F) is a strong reflection of the ontogeny of *P. pacificus*.

Further categorisation of pyrophosphatase-treated small RNAs (Fig. 1C–F) as miRNAs or piRNAs was restrained to reads between 19 and 26 nucleotides (Fig. S1A–D). Sequences classified as piRNAs were identified by the presence of the conserved Ruby motif upstream of the sequences [25, 42]. Similar to previous results [40, 41], piRNAs in *P. pacificus* have a bias for a length of 21 nucleotides. Within our analysis, relative abundance of piRNAs was lower in J2 and J3 stages than in J4/YA and embryos, correlating with the proliferation of the germline in *P. pacificus* and parental inheritance of piRNAs in *C. elegans* embryos [108, 109]. Furthermore, those small RNA sequences that had a perfect sequence match to existing mature sequences on miRBase (release 22.1) [83, 110, 111] were annotated as miRNAs. The observed increase in miRNAs of lengths 23 and 24 nucleotides during J2 and J3 was notably different from adults [41]. Lastly, the remaining reads that did not categorise as either piRNAs or miRNAs were annotated as ‘others’, which would include siRNAs. To analyse differences between expression of small RNAs categorically between different developmental stages, we performed differential expression analysis for small RNAs, resulting in a compendium of differentially expressed small RNAs (DEsRNAs) across six pairwise comparisons (Tables S31–S36). The comparison between embryos and J4/YA stages had the most DEsRNAs at 31419, whereas that between J2 and J3 stages had the least number of DEsRNAs at 3213. A detailed summary on DEsRNAs, including their categories, is also provided (Table S37).

Taken together, these results present the first comprehensive dataset of developmental small RNA transcriptome for *P. pacificus*.

### A highly conserved miRNA is the most abundant juvenile-specific miRNA in *P. pacificus*

Two developmental patterns in the small RNA profiles stood out: the large relative abundance of category 22U in the embryo with progressive reduction, and the dominance of category 24A in juveniles and young adults in contrast to embryos. To explore these further, unique reads constituting more than 1% of the total abundance of 22U at embryonic stage were identified, showing that a single sequence made up for nearly half of 22U reads (Fig. S2A), corresponding to ∼10% of the entire embryonic small RNA transcriptome. Cross-referencing with miRBase revealed that the sequence corresponded to ppc-miR-2235a-3p, a *P. pacificus* miRNA that shows a seed sequence conservation [89, 112] with the miR-35 family of *C. elegans*, a miRNA family with embryonic expression [113, 114]. Please note that in this study, we define a miRNA family based on the sharing of the same seed sequence, which does not necessarily imply shared ancestry [70]. Our definition differs from MirGeneDB’s definition of a family of miRNAs, which explicitly tries to group only homologous miRNAs into a family [115]. However, the use of seed-based definition of miRNA family is more in line with previous *C. elegans* literature. *Ppa-mir-2235a-3p* has recently been found to exist in 44 copies in the genome of the RSC011 strain of *P. pacificus*, where it has been shown to play a key role in the transgenerational inheritance of the predatory eurystomatous mouth morph, with the miRNA being destabilised by the ubiquitin ligase *Ppa-ebax-1* [116]. Interestingly, in our wild-type PS312 strain, *Ppa-mir-2235a-3p* was found to exist in 35 copies on chromosome II in an ∼8000 base-pair window, with any two copies of the 22-nucleotide miRNA-coding gene separated by a precise 202 base-pairs of consensus genomic sequence (except in between copies #18 and #19) (Fig. S2B), indicating that the arrangement of the *Ppa-mir-2235a-3p* locus seems to have undergone recent evolutionary changes. Furthermore, this genomic arrangement of *Ppa-mir-2235a-3p* is reminiscent of *mir430* cluster in zebrafish, wherein a triplet of three miRNA-coding genes — *mir430a*, *mir430b*, and *mir430c* — forms a cluster of nearly 90 copies in a 120-kb genomic locus [117]. miR-430 too is expressed early in zebrafish development, and is responsible for the deadenylation and subsequent degradation of maternally-provided mRNAs at the onset of zygotic transcription [118].

Similar to 22U, a single unique read made up the bulk of the 24A category, constituting between 60% and 70% of 24A in the three post-embryonic time points (Fig. 2A–C), and thus amounting between 15% and 20% of the entire small RNA transcriptome. This corresponds to approximately 1.5 times the abundance of ppc-miR-2235a-3p at embryonic stage. Interestingly, we could not find this small RNA read in the list of known *P. pacificus* miRNAs on miRBase. To test for the possibility of sequencing artefacts, a putative biogenesis site for the read was searched for by perfect sequence complementarity within the *P. pacificus* genome. The 24 base-pair region between the genomic coordinates 2633305 and 2633328 on chromosome X of *P. pacificus* [57] was a perfect and unique match against our query sequence. Next, leveraging the availability of sequenced genomes of other species in the family Diplogastridae [119–122], the 24-nucleotide read was queried against and gave perfect matches in eight of the 10 genomes. The two exceptions were *Pristionchus fissidentatus*, where the match was limited to the first 23 nucleotides, and *Pristionchus mayeri*, where no match was found. Furthermore, in *Pristionchus exspectatus*, the gonochoristic sister species to *P. pacificus* [122], the query sequence matched on the same chromosome as in *P. pacificus*, and in *Allodiplogaster sudhausi*, a diplogastrid nematode with a whole genome duplication [121], two perfect matches against the genome were identified. These results, coupled with the small RNA sequencing results from *P. pacificus* across multiple replicates, and the abundance of the read, strongly suggest that the 24A sequence element is not an artefact, but a bona fide and abundant small RNA with putative conservation of expression in other diplogastrid nematodes.

There was no Ruby motif upstream of this small RNA, suggesting it is not a piRNA [25, 42]. We did, however, discover complete sequence identity (taking into account, the first 22 nucleotides of the read or more) in other organisms on miRBase, corresponding to miRNA miR-100. Therefore, we named this highly abundant 24A small RNA read of *P. pacificus* as ppc-miR-100. mir-100 has been implicated in differentiation, cell signalling, and multiple types of cancer [123–131]. However, its biology in invertebrates remains largely enigmatic. Notably, although miR-100 is a highly conserved regulatory RNA with a strong sequence conservation that may predate bilaterian evolution [132], the *C. elegans* miRNA repertoire does not contain a cel-miR-100. Instead, it transcribes six miRNAs, cel-miR-51, −52, −53, −54, −55, and −56, all of which share the seed sequence with miR-100 from other organisms, but show divergence from an otherwise deeply conserved sequence (Fig. 2D) [83, 114, 132–134]. The transcription of miR-51 family in *C. elegans* is switched on from early embryonic stages. Furthermore, miR-51 family has recently been shown to be one of the two miRNA families (the other being miR-35 family) whose transcription is necessary and sufficient for embryogenesis in *C. elegans* [114], and is responsible for pharyngeal attachment in the embryo [134]. However, the sequence similarity between the cel-miR-51 family members and ppc-miR-100 ceases to exist beyond the seed region. Thus, the presence of miR-100 in *Pristionchus* nematodes provides a new and potentially powerful avenue to study its biology.

The absence of a miRNA as highly abundant and deeply conserved as ppc-miR-100 from the list of known *P. pacificus* miRNAs was intriguing. To address this conundrum further, we performed an analysis to identify novel miRNAs across the different developmental time points using miRDeep2. We also performed the same analysis on a previously published study [41], in which *P. pacificus* small RNAs were characterised in adults and dauers with and without pyrophosphatase treatment. miRDeep2 results of novel miRNA identification analysis from that previous study also identified ppc-miR-100 in its list of novel miRNAs (Table S5), as did the results for our developmental stage-specific samples, indicating that presumably, the list of novel miRNAs identified from that study have not been updated within the list of *P. pacificus* miRNAs on miRBase. Considering the novel miRNAs from that previous study as already known, we identified a total of 64 novel miRNAs across the four time points in our study (Table S4, Fig. S1E). Nearly two-thirds (40 out of 64) of these novel miRNAs were found to be expressed in only one of the four developmental time points (Fig. S1E). Relative abundance for these novel miRNAs was also low, with only two miRNAs — UGGCAAGAACAGUGGCACGGUCG and UCUUCUGUGUGCAACCCCCAAC — having notable levels of expression (approximately 0.34% and 0.16% respectively; Fig. S1F–I).

### The *let-7-complex* shows two alternative arrangements in nematodes

In *Drosophila melanogaster* and other animals, including vertebrates, the miRNA-coding gene *mir-100* occurs in a clustered arrangement with two other miRNA-coding genes, *let-7* and *mir-125* [69, 70, 135] (Fig. 3A). miR-100 and miR-125 are both members of an ancient lineage of miRNAs that has been called the MIR-10 family in MirGeneDB [70, 136]. The *D. melanogaster* cluster locus is polycistronic, giving rise to the precursors of the three miRNAs from the same primary transcript, and is known as the *let-7-complex* [14, 137]. *C. elegans* is an exception to this arrangement. The orthologues of both *let-7* and *mir-125* (the latter called *lin-4*) are located on different chromosomes (Fig. 3B) [135]. Both *lin-4* and *let-7* are important for the temporal control of developmental decision-making in *C. elegans* [10, 12], albeit at different stages of the worm’s ontogeny. This correlates with the genomically separate arrangement of the two miRNA-coding genes in *C. elegans* on different chromosomes (*Cel-let-7* is on chromosome X and *Cel-lin-4* is on chromosome II), thus untying the polycistronic, simultaneous expression of the *let-7-complex* observed in *D. melanogaster* [135]. As for *mir-100* orthologues, while three out of six genes (*Cel-mir-54, Cel-mir-55, Cel-mir-56*) coding for miR-51 family in *C. elegans* are found on chromosome X, same as *Cel-let-7*, and form a cluster of their own [134], the chromosomal distance between the two is approximately 1.5 megabases, and thus not linked.

To determine if the three miRNA-coding gene constituents of *let-7-complex* in *P. pacificus* constitute a locus similar to the one in *D. melanogaster* and other animals (ppc-let-7 and ppc-lin-4 have already been identified in previous studies and are annotated in miRBase), we next studied their genomic arrangement. We found that both *Ppa-lin-4* and *Ppa-let-7* reside on the same chromosomes as their *C. elegans* counterparts. Interestingly, *Ppa-mir-100* and *Ppa-let-7* are tightly linked, with *Ppa-let-7* being only 200 base-pairs downstream and in the same orientation as *Ppa-mir-100* (Fig. 3C), reminiscent of the arrangement in the *let-7-complex*. Exploring the genomic arrangements of these three miRNA-coding genes in other nematodes, we found that *Caenorhabditis briggsae*, *Caenorhabditis remanei*, and *Caenorhabditis brenneri* all follow the same pattern as *C. elegans* in not having a ‘classical’ *mir-100* within their genome, and having *lin-4* and *let-7* genes dispersed over different chromosomes or genome contigs (Fig. 3B). Conversely, four nematode species with *mir-100* within their genomes — *Ascaris suum*, *Panagrellus redivivus*, *Heligmosomoides polygyrus*, and *Brugia malayi* — all have a *P. pacificus*-like arrangement in which *mir-100* and *let-7* are tightly linked in a ∼200 base-pair region, whereas *lin-4* is found on a different chromosome or contig (Fig. 3C). These four species and *P. pacificus* are spread over three nematode clades (two are free-living, three are parasitic) [138]. This suggests a progressive disintegration of the *let-7-complex* in nematodes, whereby the most downstream miRNA-coding gene of the three within the locus, *mir-125*/*lin-4*, was the first to dissociate from the rest of the locus, as it became relocated to a different chromosome in the last common ancestor of all nematodes investigated in this study. This was followed by the loss of the most upstream miRNA-coding gene, seen here by the absence of *mir-100* orthologues from the locus in all investigated *Caenorhabditis* species. In a final step, *let-7* itself may get lost, as has been reported recently in multiple species within the *japonica* group of *Caenorhabditis* [139].

### The expansion of miR-100 family precedes the loss of the *let-7-complex* in *Caenorhabditis* nematodes

Despite the sequence similarity in the seed regions between the miR-51, −52, −53, −54, −55, and −56 of *C. elegans* and miR-100 from *P. pacificus*, *A. suum*, *D. melanogaster*, and *H. sapiens* (Fig. 2D), it is unclear whether the two sets of miRNAs are orthologous in the sense of shared ancestry. While miR-100 may have diverged and expanded into the miRNAs miR-51, −52, −53, −54, −55, and −56 in the *C. elegans* lineage, theoretically, it is also possible that miR-100 got replaced by a non-orthologous miRNA that had evolved the same seed sequence as miR-100 independently, which would constitute a case of non-orthologous gene displacement at the level of miRNAs [140]. In support for this, miR-51, −52, and −53 are derived from the 5’ arm of the precursor, whereas miR-54, −55, and −56 are derived from the 3’ arm. This has been interpreted as strong evidence supporting convergent evolution of miRNAs with identical seeds [70]. Thus, miRNAs cannot be considered as homologous just because they share the same seed sequence. In an attempt to trace the evolutionary relationships among members of miR-100 family, we performed a phylogenetic analysis of all miR-100 family members (Fig. S9A). Although the limited number of phylogenetically informative positions in a multiple sequence alignment of miRNAs sharing the same sequence does not allow a conclusive interpretation of the evolutionary history, it shows that *P. pacificus* has 12 members of the miR-100 family — distributed across *P. pacificus* chromosomes I, IV, and X — of which some may be potentially orthologous to miRNAs of the miR-54–56 cluster in *C. elegans* (Fig. S9A) [134]. The nematode *A. suum* also seems to have an orthologue of miR-56 of *C. elegans*, consistent with the grouping of several of these sequences into the same family by MirGeneDB. This suggests that the expansion of miR-100 family occurred before the loss of the *let-7-complex* in *Caenorhabditis* nematodes. Additionally, this excludes miRNAs miR-54, −55, and −56 of *C. elegans* as potential orthologues of miR-100. The remaining *C. elegans* miR-100 family members — miR-51, −52, and −53 — appear to have orthologues only in other *Caenorhabditis* species, which is consistent with them being miR-100 orthologue candidates. The phylogenetic tree also shows the miR-100 orthologue in the strongylid *H. polygyrus*, which is even more closely related to *C. elegans* than *P. pacificus*. This assigns the period of miR-100 transition close to the ancestor of the *Caenorhabditis* genus. As the overall topology of the phylogenetic tree is poorly supported, we complemented the phylogenetic analysis by visualisation of differences (gaps or substitutions) in the multiple sequence alignment as a heat map (Fig. S9B). This shows that every miRNA that is annotated as miR-100 is less than five differences away from another miRNA annotated as miR-100 in another species. Based on the overall similarity pattern, we arbitrarily grouped the majority of miR-100 family members into one of the following four groups: G51, G52, G55, and G100. G100 corresponds to most miR-100 orthologues from diverse animals, whereas G51, G52, and G55 are named according to miRNAs miR-51, −52, and −55 of *C. elegans* that are respectively members of these groups. Both *Caenorhabditis*-specific groups G51 and G52 exhibit some similarity with members of G100, which supports potential orthology between *C. elegans* miRNAs miR-51, −52, and −53 (miR-53 is a member of G52), and miR-100 from other species. Based on these data, we would propose that in an ancestral *Caenorhabditis* lineage, miR-100 evolved into miR-51, −52, and −53, with miR-51 being the best candidate as the orthologue of miR-100 in *C. elegans* (Fig. S9C). This is also supported by the grouping of miR-51 and miR-100 into the MIR-10 family by MirGeneDB [70]. However, given the extensive sequence divergence between miR-51 and miR-100 orthologues from other species, we cannot conclusively decide whether these miRNAs are truly orthologous in the sense of shared ancestry or whether miR-100 got replaced by non-orthologous miRNAs. At the level of miRNA function, genetic mutants for miRNA-coding genes *mir-51*, *mir-52*, *mir-53*, *mir-54*, *mir-55*, and *mir-56* in *C. elegans* — of individual genes as well as in various combinations — result in pleiotropic developmental defects, with the phenotypic intensity of defects varying according to the combination of genes perturbed [133, 134, 141]. The strongest defect, resulting in lethality, was reported when at least five miRNA-coding genes (all except *mir-53*) out of the six were mutated [133, 134, 141]. A recent study [141] systematically characterising the function of these six miRNAs in *C. elegans* reported pleiotropic phenotypes, including reduction in brood size, uterine egg retention, abnormal egg-laying, and locomotory defects, due to disruption of extracellular matrix-dependent signalling pathways, in line with our observations and analyses in the context of miR-100 in *P. pacificus*. However, functional differences amongst the three miRNAs miR-51, −52, and −53 of *C. elegans* in comparison to miR-100 of *P. pacificus* in terms of orthologous direct mRNA targets are wide [134, 141] (please see our sub-sections ‘Phenotypic characterisation of *P. pacificus* miRNA mutants’ and ‘miR-100 and let-7 regulate extracellular matrix-regulating developmental genes in *P. pacificus*’ in this manuscript, below).

### Phenotypic characterisation of *P. pacificus* miRNA mutants

To characterise the components of the *let-7-complex* in *P. pacificus* further, we investigated the expression levels of the three miRNAs (Fig. 3D–F). As reported above, miR-100 was found to be highly abundant during J2 and J3 stages, accounting for ∼20% of the entire small RNA transcriptome at its peak. Similar to miR-100, let-7 also peaked at J3, indicating the possibility that their origins might be from a polycistronic locus as in *D. melanogaster* despite the differences in their relative abundance levels [142]. Finally, lin-4 was the least abundant of the three, peaking at nearly 0.2% of the total during J3. The maxima of abundance attained by the three miRNAs at the same developmental stage led us to explore their putative roles in *P. pacificus*.

We used CRISPR/Cas9-mediated genome editing [56, 58] to generate mutants of *Ppa-mir-100*, isolating seven independent alleles (Fig. S7A). First and foremost, all homozygous *Ppa-mir-100* mutants were viable. However, *Ppa-mir-100* mutants were variably sick, showed egg retention, moulting defects, slower locomotion, and a mild him (<u>h</u>igh incidence of <u>m</u>ales) phenotype. To quantify these differences further, we used synchronised egg clutches to measure egg-laying rates, progeny survival, and overall brood sizes, eliminating potential effects of maternal age (Fig. S4; see also ‘Phenotypic assays’ in the Methods section for details). We found that *Ppa-mir-100* mutants showed an overall reduction in egg-laying rate, albeit with varying severity (Fig. S3A). Using *Ppa-mir-100(tu1684)* for further phenotypic characterisation, we scored for embryonic lethality and survival to adulthood, in addition to egg-laying rate. *Ppa-mir-100(tu1684)* mutants performed worse than wild-type, *i.e.*, had higher embryonic lethality, lower survival to adulthood, and lower egg-laying rate, than wild-type (Fig. 4C–E). Overall, brood sizes of *Ppa-mir-100(tu1684)* worms were noticeably lower than those of wild-type over the course of five days (Fig. 4A). Resolving the lattermost data temporally showed that *Ppa-mir-100(tu1684)* mutants laid fewer eggs at the peak of reproductive activity, and maintained their production for a longer window than wild-type (Fig. 4B).

Next, we generated *Ppa-let-7* mutants. We obtained 11 mutations (Fig. S7B), which were also not lethal under lab conditions, but showed stronger pleiotropic phenotypes than those observed in *Ppa-mir-100* mutants (sick worms, bagging of eggs in the uterus, moulting defects). Additionally, we observed larval arrest, mid-body rupture, internal hatching, dumpy, and blistered worms. Phenotypic characterisation showed that disparities in egg-laying rates were stronger and more consistent, with statistically significant differences across all 11 alleles (Fig. S3B). We observed that *Ppa-let-7(tu1719)* mutants had lower egg-laying and survival-to-adulthood rates than *Ppa-mir-100(tu1684)* mutants, but showed marginally lower embryonic lethality than the latter, although still higher than wild-type (Fig. 4C–E). *Ppa-let-7(tu1719)* mutants also had smaller overall brood sizes than *Ppa-mir-100(tu1684)* mutants (Fig. 4A–B).

To test if a double mutant of both miRNAs would have a stronger phenotype, we generated *Ppa-mir-100* mutants in *Ppa-let-7(tu1724)* background, yielding four *Ppa-mir-100 Ppa-let-7(tu1724)* double mutants (Fig. S7C). These mutants too were viable, and had significantly lower egg-laying rates than wild-type worms (Fig. S3C), but did not show a strong enhancement of the pleiotropic phenotype of the two single mutants. Using *Ppa-mir-100(tu1898) Ppa-let-7(tu1724)* for subsequent phenotypic analyses showed that the double mutant had a stronger phenotype than *Ppa-mir-100(tu1684),* but weaker than *Ppa-let-7(tu1719)* mutants (Fig. 4A–E).

Lastly, we generated *Ppa-lin-4* mutants and obtained two independent alleles (Fig. S7D). Surprisingly, *Ppa-lin-4(tu1943)* mutants showed no noticeable differences when compared to wild-type worms when scoring for their egg-laying rate, embryonic lethality, and survival to adulthood (Fig. 4C–E). In a similar vein, the brood size of *Ppa-lin-4(tu1943)* mutants was not noticeably different from that of the wild-type worms, although admittedly, with a higher variation among individual samples (Fig. 4A). Interestingly, the difference lay in the temporal distribution of the brood size, wherein the bulk of the *Ppa-lin-4(tu1943)* progeny were laid more than a day later than the wild-type worms (Fig. 4B), indicating towards an aberrant but viable temporal developmental shift. The original *lin-4* mutant strain (e912) in *C. elegans* was investigated by the Ruvkun and Ambros labs in the 1990s because of its striking developmental timing defects, wherein *lin-4* mutants exhibited heterochronic defects by extension of juvenile cells and moulting phenotypes into adults [10, 143]. Our results on the mutant *lin-4* phenotype in *P. pacificus* open new avenues to explore the biology of this important miRNA in comparative evolutionary and developmental contexts.

### Small RNA sequencing of mutants confirms absence of miRNAs and suggests partial phenotypic rescue by other miR-100 family members in *P. pacificus*

To study the small RNA profiles in miRNA mutants, J3 stage-specific small RNA sequencing of wild-type, *Ppa-mir-100(tu1684)*, *Ppa-let-7(tu1719)*, and *Ppa-mir-100(tu1898) Ppa-let-7(tu1724)* was performed (3 biological replicates for *Ppa-mir-100(tu1684)*, 2 for the rest). The J3 stage was selected since both miR-100 and let-7 were the most abundant at this stage (Fig. 3D–E). Small RNA profiles showed that 24A and 22U, housing miR-100 and let-7, respectively, had lower relative abundances in *Ppa-mir-100(tu1684)* and *Ppa-let-7(tu1719)* mutant worms, respectively, in comparison to wild-type, and both were less abundant in the *Ppa-mir-100(tu1898) Ppa-let-7(tu1724)* double mutant (Fig. 5A–D). Indeed, we found that ppc-miR-100 and ppc-let-7 reads were completely absent in the corresponding mutants (Fig. 5M–N). This result supports that the generated mutants represent knockouts of these miRNAs.

Frequently, individual miRNA mutants show weak to no phenotypes, due to functional redundancy provided by other miRNAs that belong to the same family [133, 134, 144–146]. To test the possibility that the variable phenotypes across *Ppa-mir-100* mutants could be due to redundancy with other miR-100 family members being potentially transcribed in the *P. pacificus* genome and resulting in partial stochastic rescue of the phenotypes across alleles, we queried for the expression levels of entire miR-100 family in wild-type and mutant worms. Majority of the miR-100 family in wild-type worms was constituted by miR-100 itself, which was abrogated in the *Ppa-mir-100(tu1684)* and *Ppa-mir-100(tu1898) Ppa-let-7(tu1724)* double mutants, but not in *Ppa-let-7(tu1719)* mutants (Fig. 5E–H). However, a small number of 22 nucleotide-length reads belonging to the miR-100 family were also found in wild-type worms (Fig. 5E), which were still present in *Ppa-mir-100(tu1684)* and the *Ppa-mir-100(tu1898) Ppa-let-7(tu1724)* double mutants (Fig. 5F, H), indicating that these are miR-100 family members which could potentially partially rescue the loss of miR-100, thus resulting in an incomplete and inconsistent mutant phenotype (Fig. S3A) [134, 144, 145]. Such a pattern was not seen for let-7; the expression of the let-7 family in *P. pacificus* is primarily composed of let-7 itself, which, when knocked out, has nearly no family members being expressed to rescue its loss (Fig. 5I–L), resulting in a stronger and more consistent phenotype (Fig. S3B).

Next, to rule out the possibility that miR-100 itself might be partially present in the alleles with weaker phenotypes and may have only abrogated in the phenotypically strongest allele, four more mutant alleles across the three mutants — *Ppa-mir-100(tu1638)*, *Ppa-mir-100(tu1685)*, *Ppa-let-7(tu1724)*, and *Ppa-mir-100(tu1900) Ppa-let-7(tu1724)* double mutant — were sequenced (3 biological replicates for *Ppa-mir-100(tu1638)* and *Ppa-mir-100(tu1685)*, 2 biological replicates for *Ppa-let-7(tu1724)* and *Ppa-mir-100(tu1900) Ppa-let-7(tu1724)* double mutant), and then searched for presence of miR-100 and let-7. Regardless of their phenotypes, or lack thereof, no miR-100 or let-7 was found in corresponding mutants (Fig. S5A–B). Furthermore, our analysis of the expression levels of entire miR-100 family in *Ppa-mir-100(tu1638)* and *Ppa-mir-100(tu1685)* alleles, and of the let-7 family in *Ppa-let-7(tu1724)* allele (Fig. S5D–F) reciprocated our previous results for *Ppa-mir-100(tu1684)* and *Ppa-let-7(tu1719)* respectively (Fig. 5F, K); the two *mir-100* alleles showed expression of other miR-100 family members, but the *let-7* allele showed no expression of other let-7 family members. Thus, the presence of other miR-100 family members, and not partial presence of miR-100 itself in the phenotypically weak alleles, is likely responsible for incomplete phenotypic penetration in *Ppa-mir-100* alleles.

Lastly, we performed a PCA of our small RNA transcriptomes for all replicates of the three mutant genotypes and wild-type (19 samples in total), which revealed that the small RNA transcriptomes of all *Ppa-mir-100* mutants form a distinct cluster (Fig. S5C). This result indicates that irrespective of the phenotypic variation observed across different mutant alleles, the effect of mutation of *Ppa-mir-100* on the small RNA profiles is consistent across all tested alleles, and is therefore gene-specific. Additionally, we also performed differential expression analysis of small RNAs between the wild-type and the three *Ppa-mir-100* alleles (Tables S38–S43). The results across the six pairwise comparisons revealed a difference of at least an order of magnitude between the number of DEsRNAs in comparisons between the wild-type genotype and either of the *Ppa-mir-100* alleles (PS312 *vs. Ppa-mir-100(tu1638)*: 5504 DEsRNAs; PS312 *vs. Ppa-mir-100(tu1684)*: 6087

DEsRNAs; PS312 *vs. Ppa-mir-100(tu1685)*: 5166 DEsRNAs) versus comparisons among the three *Ppa-mir-100* alleles (*Ppa-mir-100(tu1684) vs. Ppa-mir-100(tu1638)*: 645 DEsRNAs; *Ppa-mir-100(tu1684) vs. Ppa-mir-100(tu1685)*: 41 DEsRNAs; *Ppa-mir-100(tu1638) vs. Ppa-mir-100(tu1685)*: 509 DEsRNAs), complementing our PCA results that the three *Ppa-mir-100* alleles are less indistinguishable among themselves then when compared to the wild-type genotype. However, we cannot completely rule out a contribution of CRISPR-induced off-target effects towards the observed phenotypic variation, and as such, elucidating the exact role of miR-100 and mechanism of partial phenotypic rescue by miR-100 family members requires further experimental characterisation.

### miR-100 and let-7 regulate extracellular matrix-regulating developmental genes in *P. pacificus*

We identified putative direct mRNA targets of miRNAs miR-100 and let-7 through perfect complementarity to the seed region (taking into account nucleotides 2–7, instead of 2–8, to procure a more inclusive list of targets) of the miRNAs within the presumptive 3’ UTRs of gene annotations of the *P. pacificus* genome [92]. We identified 1095 and 2951 potential target mRNAs with regions complementary to the seed regions of miR-100 and let-7 respectively (Tables S6–S7). From some of the known targets of let-7 in *C. elegans* [12, 147–149], we found the orthologues of *Cel-hbl-1*, *Cel-daf-12*, and *Cel-let-60* in the *P. pacificus* genome — PPA26451, ppa_stranded_DN30554_c0_g2_i14, and PPA03306 respectively — among the putative targets of let-7 in *P. pacificus*, but did not find those of *Cel-lin-28* (PPA39985) and *Cel-lin-41* (ppa_stranded_DN18733_c0_g1_i1) amongst the list of putative let-7 targets (Table S7). For miR-100, however, the overlap was sparser, with only the *Cel-kal-1* orthologue (ppa_stranded_DN30965_c9_g5_i2) identified as a putative direct target of miR-100 in *P. pacificus*, whereas those of *Cel-hst-3.2* (PPA32231), *Cel-hst-6* (PPA28737), *Cel-hst-1* (ppa_stranded_DN31306_c0_g3_i8), *Cel-lin-12* (ppa_stranded_DN31313_c0_g1_i1/ppa_stranded_DN31313_c0_g1_i2), *Cel-mig-1* (PPA37134), *Cel-lin-17* (Iso_D.13440.3), *Cel-lon-1* (PPA34110), *Cel-vab-2* (PPA28315), *Cel-efn-2* (PPA05071), *Cel-efn-4* (PPA14865), and *Cel-cdh-3* (ppa_stranded_DN31465_c0_g2_i1) were not (Table S6) [134, 141], pointing towards only partial conservation among the targets of the orthologous miRNAs between the two species.

To identify putative genes regulated by the *P. pacificus* miRNAs, we performed RNA sequencing of mutant strains (3 biological replicates for *Ppa-mir-100(tu1684)*, 2 for the rest). *Ppa-mir-100(tu1684)* mutant worms had the most differentially expressed genes (DEGs), with a total of 548, a majority of which (449 genes) were upregulated in comparison to wild-type, while the remaining 99 genes were downregulated (Fig. 6A, Table S1). *Ppa-let-7(tu1719)* mutants had fewer DEGs, with a total of 224, of which approximately one half was upregulated (116 genes) and the other (108 genes) downregulated (Fig. 6B, Table S2). Lastly, the *Ppa-mir-100(tu1898) Ppa-let-7(tu1724)* double mutant had a total of 214 DEGs, with 31 downregulated and 183 upregulated genes (Fig. 6C, Table S3).

The skewed distribution of upregulated versus downregulated DEGs in the *Ppa-mir-100(tu1684)* and *Ppa-mir-100(tu1898) Ppa-let-7(tu1724)* double mutants was intriguing (Fig. 6A–C). Therefore, we looked at the overlap of DEGs between the three mutants in a pairwise manner, and then across all three together. A total of 67 genes were commonly differentially expressed between the two single mutants, whereas *Ppa-mir-100(tu1684)* mutant and the double mutant shared 85 DEGs, and *Ppa-let-7(tu1719)* mutant and the double mutant shared 71 DEGs. Also, there were 35 DEGs shared among all three mutants (Fig. 6D). Looking at the direction of differential expression in each pairwise comparison, the number of common genes oppositely regulated between the double mutant and the single mutants was 2/85 and 4/71 for *Ppa-mir-100(tu1684)* and *Ppa-let-7(tu1719),* respectively. The common genes between the two single mutants stood apart from the rest: 14/67 common genes between *Ppa-mir-100(tu1684)* and *Ppa-let-7(tu1719)* mutants were oppositely regulated, none of which were present in the common genes among all three mutants. In *Ppa-let-7(tu1719)* mutants, 11 of these 14 were downregulated, with 10 out of those 11 strongly downregulated (log_2_(fold change) < −2). Thus, let-7, in parts, oppositely regulates genes regulated by miR-100, an effect that is abrogated in the double mutant, which we hypothesise explains the more modest effect in its phenotype.

Comparing the DEGs with putative direct mRNA targets of miRNAs, we found that 19 out of the 449 upregulated genes (4.23%) and 2 out of the 99 downregulated genes (2.02%) in *Ppa-mir-100(tu1684)* were also predicted to be direct targets of miR-100. For let-7, we found 10 out of 116 upregulated genes (8.62%) and 13 out of 108 downregulated genes (12.04%) as predicted targets in *Ppa-let-7(tu1719)* mutants. These admittedly small overlaps may have several explanations. First, RNA-seq has limited utility in investigating miRNA targets, since miRNAs can regulate translation of mRNA targets without necessarily or strongly affecting mRNA abundance. Moreover, in cases where a target mRNA may be broadly expressed but regulated by the miRNA in only a subset of cells, removing the latter may not result in a detectable change in the abundance of the former in extracts of whole animals. Second, our prediction pipeline only searched for perfect seed matches in the 3’ UTR regions of annotated genes, and imperfect matches in the 3’ UTR or matches in other parts of the mRNA were ignored. Third, it may be possible that most of the observed differential expression is due to indirect effects downstream of the direct mRNA targets of the two miRNAs.

To explore the DEGs further, we performed an enrichment analysis of their protein domains, identifying domains with an enrichment of 1 or higher, and their corresponding Rich Factors (Fig. 6E–J). For each mutant, upregulated genes had stronger statistical support by the data for enrichment (*i.e.*, smaller *P*-values) and more domains with high degree of enrichment (*i.e.*, higher Rich Factors) than their downregulated counterparts. *Ppa-mir-100(tu1684)* mutant worms showed the strongest support within the data for collagen and nematode cuticle domains in both upregulated and downregulated genes, indicating a general perturbation of these structural proteins. For *Ppa-let-7(tu1719)* mutants, the statistical support for collagen and nematode cuticle-containing proteins in the upregulated genes was accompanied by a strong support for astacins in the downregulated genes (Fig. 6G). Astacins are a class of developmental genes highly conserved across animals, and encode for metalloproteases, a conserved protein family known for their zinc-dependent extracellular proteolytic activity during development in multiple organisms [150]. Humans have six genes encoding for astacins, which includes BMP1 (bone morphogenetic protein 1), whereas invertebrates such as *C. elegans* have nearly 40 [150–152]. In mice and sea urchins, astacins are responsible for the cleavage of a protein-rich layer of vitelline envelope (zona pellucida in higher animals) to facilitate hatching and to prevent polyspermy post fertilisation [153, 154]. In the context of nematodes, astacins are known to participate in multiple developmental pathways. The first astacin identified and characterised in *C. elegans* was *hch-1*. HCH-1 is an embryonic protein, whose absence leads to a delayed albeit viable hatching in *C. elegans* [155]. However, the most common pathway that nematode astacins are known to play a role in is moulting of its cuticle. *nas-37*, *nas-36* and *dpy-31*/*nas-35* were found to be expressed in the hypodermal cells, and mutants for these genes were found to have moulting defects [156–158] due to incomplete degradation of the older cuticle, an extracellular matrix layer composed of collagen [159, 160]. Astacins are capable of regulating collagen levels by processing of procollagen [161], and subsequent research has shown that moulting-related functions of astacins are evolutionarily conserved between free-living and parasitic nematodes [162–165].

Domain enrichment analysis for the *Ppa-mir-100(tu1898) Ppa-let-7(tu1724)* double mutant broadly recapitulated the results of *Ppa-let-7* worms; astacin was statistically the most supported domain by the data in downregulated genes, whereas collagen and cuticle were the most statistically supported domains within the upregulated genes (Fig. 6I–J). There was also an overall increase in the total number of domains showing enrichment in the double mutant, and a strong increase in ribosomal protein domains in the upregulated genes (Fig. 6J).

In *P. pacificus*, the interest in astacins goes beyond their presumed functional conservation in moulting, and into their potential roles in development of plastic adult mouth morphs [166, 167]. Recently, Sieriebriennikov and colleagues identified that the common regulatory targets of two nuclear hormone receptors which control feeding plasticity in *P. pacificus*, *Ppa-nhr-40* and *Ppa-nhr-1*, are enriched in astacins [166]. We found 15 out of 24 common transcriptional targets of *Ppa-nhr-1* and *Ppa-nhr-40* to be overlapping with the DEGs in *Ppa-let-7* mutants, eight of which were also overlapping with the DEGs in the double mutants (Table S9). Out of the 12 astacins that were found among the 24 NHR targets, eight were differentially expressed in *Ppa-let-7* worms, five of which were also differentially expressed in the double mutant. All of these genes were downregulated in our miRNA mutants (Table S9). The unexpected regulatory convergence on astacins and collagens may suggest that these miRNAs have a potential role in regulation of feeding plasticity too [116]. Small RNAs are known for their ability to respond to environmental perturbations and even convey the information across generations [168, 169]. Although we did not see any mouth-morph phenotypes in either of the three mutants (Eu morph > 95% in wild-type and miRNA mutants; n = 90–100 over three biological replicates), in line with the duodecuple astacin mutant [166], future analyses will reveal the extent to which small RNAs contribute to such phenotypic plasticity.

Finally, to gain biological insights about the DEGs, we performed overrepresentation analyses of our DEGs in two recently published genome-wide datasets, categorising *P. pacificus* genes on the basis of co-expression modules and identifying oscillating genes in the genome [93, 94]. Gene module overrepresentation analysis showed an overrepresentation of genes within module #11 (FDR corrected P-value < 0.001), which includes genes involved in cuticle development and maintenance, across all three mutants (Fig. S6A–C). Other modules with an overrepresentation in at least one mutant were associated with developmental pathways such as oogenesis (module #7) [108], or expression in tissues undergoing rapid developmental changes, such as the intestine (module #3) and regions associated with mouth dimorphism (module #24) (FDR corrected P-value < 0.001, Fig. S6A–C). Overrepresentation analysis within the oscillating genes showed that all three mutants had a higher representation of oscillating genes than the genome-wide baseline of ∼10% (Fig. S6D) [93]. Most of the oscillations coincide with the moulting cycles and coordinate systematic progression through developmental stages. These results also suggest that the miRNA and their target genes do not seem to be strictly co-expressed at the level of mRNAs (Fig. 3, Fig. S10). Taken together, miRNAs miR-100 and let-7 are important for overall developmental progression in *P. pacificus*, and their perturbation leads to pleiotropic defects — characteristic of developmental genes — as shown above (Fig. 4).

## Conclusions

In this study, we provide the first description of the developmental repertoire of small RNAs in *P. pacificus* (Fig. 1C–J, Fig. S1A–D), and additionally identify 64 novel miRNAs expressed in this species across different stages of its ontogeny (Fig. S1E–I). Together, these datasets can serve as a molecular resource for the nematode community to study the roles of small RNAs in an evo-devo context.

We identify that an ancient, highly conserved miRNA, miR-100, is present as the most abundant miRNA in juvenile stages of *P. pacificus* (Fig. 2A–C). Despite its transcriptional abundance, miR-100 is non-essential for viability of the worm, likely due to functional redundancy and partial rescue by other miR-100 miRNA family members (Fig. 5E–H) [133, 145]. We hypothesise that a complete knockout of the miR-100 family in *P. pacificus* would result in stronger phenotypes, possibly even lethality.

The classical *let-7-complex* is non-existent in nematodes, instead presenting its constituent miRNA-coding genes in two alternative genomic arrangements (Fig. 3B–C). This dissociation indicates a (potentially ongoing) disintegration of the *let-7-complex* in the lineages leading to *Pristionchus* and *Caenorhabditis*, where the individual components of the locus are progressively rearranged within the respective genomes. While the relocation of *lin-4* to a different chromosome appears to date back to the ancestor of all species investigated in this study, *C. elegans* shows and additional dissociation of *mir-100* from the locus along with an expansion into a family of six miRNAs with sequence divergence outside of the shared the seed sequence with miR-100 (Fig. 2D) [134]. Our phylogenetic analysis suggests that this expansion predated the loss of *let-7-complex* in the lineage leading to *Caenorhabditis* genus.

Lastly, we show that the two miRNAs — ppc-miR-100 and ppc-let-7 — as the two constituents of the remnants of *let-7-complex* in *P. pacificus*, ensure normal development by regulating transcription of collagens and astacins (along with other protein families, such as heat-shock proteins (HSPs), ribosomal protein-subunits, and the von Willebrand factor (VWF) domains, among others) (Fig. 6E–J), two developmental gene classes vastly expanded in nematodes, and important for developmental progression to adulthood.

## List of abbreviations

RNA: ribonucleic acid
miRNA: microRNA
siRNA: small interfering RNA
piRNA: PIWI-interacting RNA
C. elegans: Caenorhabditis elegans
P. pacificus: Pristionchus pacificus
RNAi: RNA interference
RdRP: RNA-dependent RNA polymerase
DNA: deoxyribonucleic acid
YA: young adult
PCA: principal component analysis
kb: kilobase
mRNA: messenger RNA
D. melanogaster: Drosophila melanogaster
C. briggsae: Caenorhabditis briggsae
C. remanei: Caenorhabditis remanei
C. brenneri: Caenorhabditis brenneri
A. suum: Ascaris suum
H. polygyrus: Heligmosomoides polygyrus
him (phenotype): high incidence of males
DEsRNAs: differentially expressed small RNAs
DEGs: differentially expressed genes
HSP: heat-shock protein
VWF: von Willebrand factor
NGM: nematode growth medium
E. coli: Escherischia coli
NGS: next generation sequencing
tRNA: transfer RNA
rRNA: ribosomal RNA
BAM: binary alignment map
crRNA: CRISPR RNA
tracrRNA: trans-acting CRISPR RNA
PAM: protospacer adjacent motif
RFP: red fluorescent protein
MANOVA: multivariate analysis of variance

**Fig. S1:**
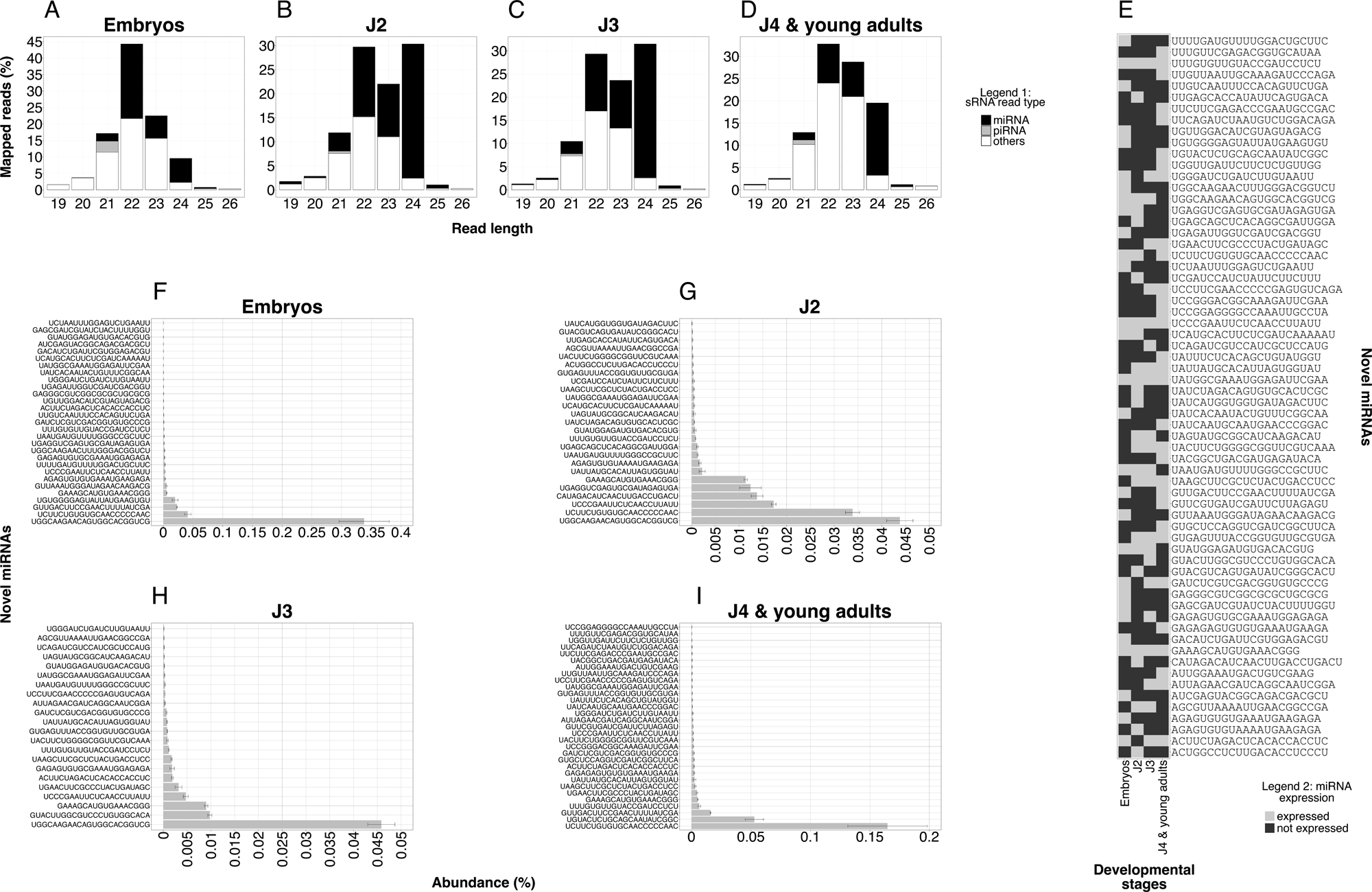
Developmental stage-specific expression of novel miRNAs of *P. pacificus*: A–D: Developmental small RNA profile, with small RNAs categorised on the basis of their length (x-axis) and small RNA type (Legend 1). Y-axis denotes the abundance (mean of three biological replicates) of each small RNA type as a fraction of the whole small RNA transcriptome for that corresponding stage. Note that the y-axis is different for different developmental stages, and also see Fig. 1C–F for comparison. E: Heat map showing expression (Legend 2) across developmental stages (x-axis) of all novel miRNAs (arranged in descending alphabetical order on y-axis) identified in this study. F–I: Developmental stage-specific abundance (x-axis; mean of three biological replicates) of novel miRNAs (arranged in increasing order of abundance on y-axis) identified in this study. Error bars represent the SEM. Note that the x-axis is different for different developmental stages.

**Fig. S2:**
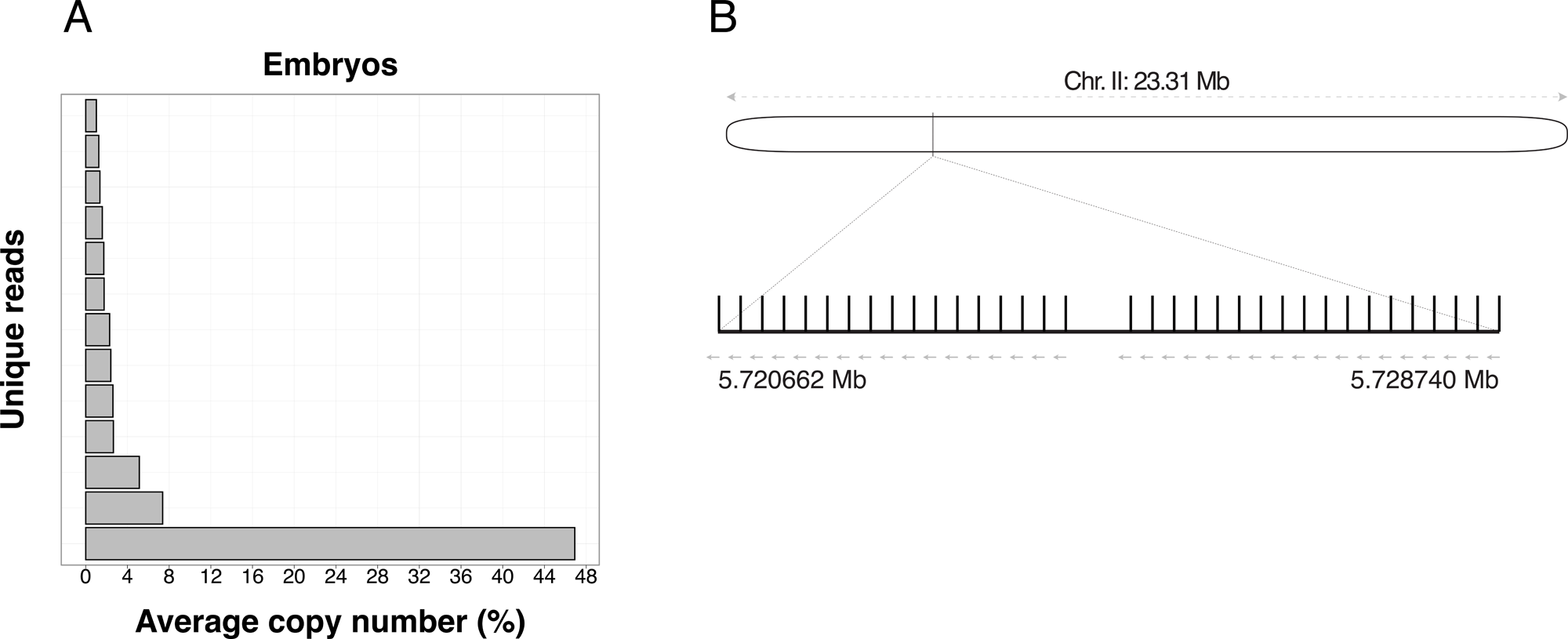
Embryonic expression level and chromosomal arrangement of the miRNA ppc-miR-2235a-3p: A: Most abundant 22U reads in embryos, represented as percentage of total 22U reads (mean of three biological replicates). The bottom-most read, with sequence UCACCGGGAGCAUUGUAUGAUC, corresponds to miRNA ppc-miR-2235a-3p. B: Schematic representation of *Ppa-mir-2235a-3p* locus on chromosome 2. The locus encompasses a length of 8079 bp, and contains 35 equidistant copies (except between copies #18 and #19) of ppc-miR-2235a-3p on the negative strand of the chromosome.

**Fig. S3:**
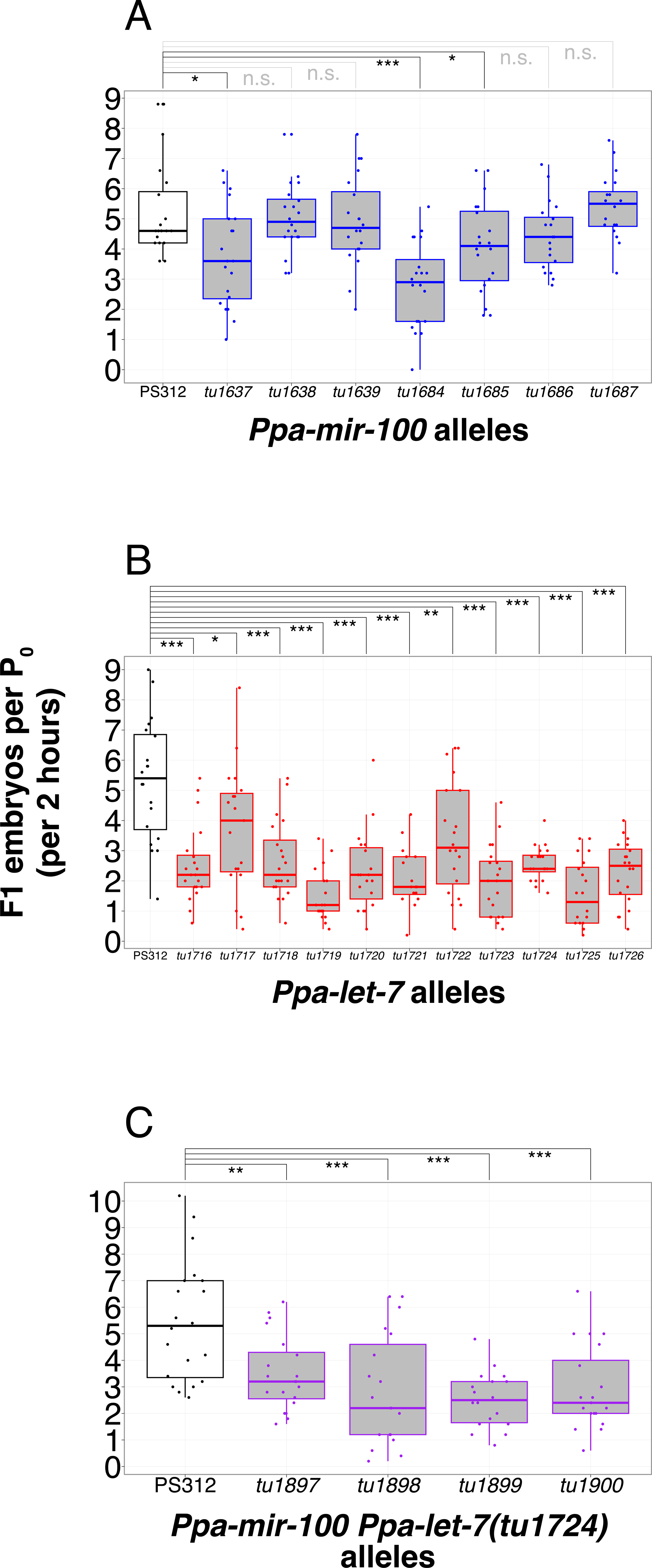
Inter-allelic phenotypic variation among *Ppa-mir-100*, *Ppa-let-7*, and *Ppa-mir-100 Ppa-let-7* double mutants: A–C: Frequency of egg-laying over 2 hours per synchronised mother for each allele of the three mutant genotypes. n = 100 mothers per allele per mutant. Please note that in the case of *Ppa-mir-100 Ppa-let-7* double mutants (C), while the four mutant alleles are all consistently and statistically significantly distinct from the wild-type genotype, the small amount of phenotypic variability within the alleles could be due to inter-allelic variation between independently generated *mir-100* mutations in the same *let-7* background.

**Fig. S4:**
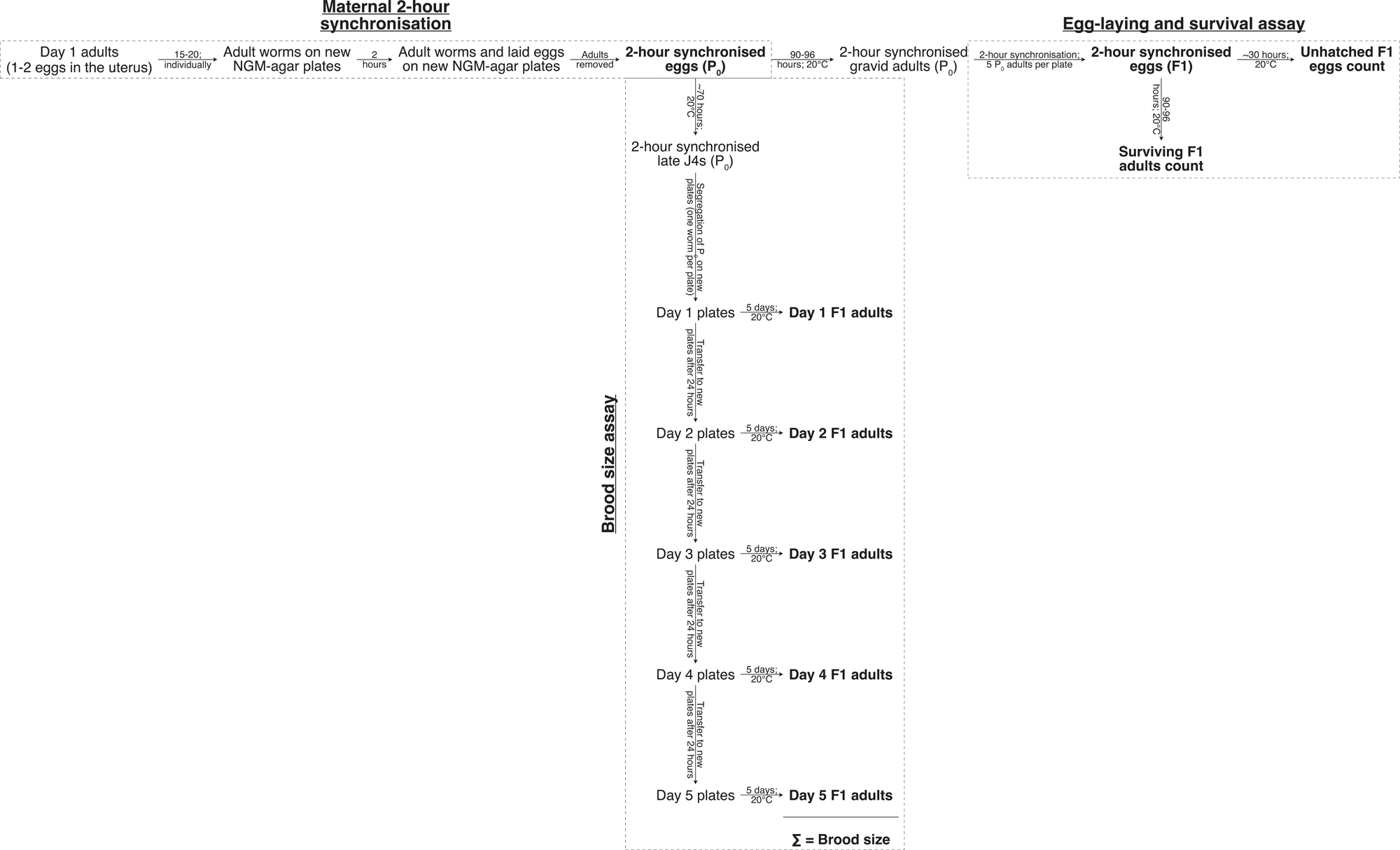
Schematic representation of phenotypic assays for *Ppa-mir-100*, *Ppa-let-7*, *Ppa-mir-100 Ppa-let-7*, and *Ppa-lin-4* mutants: Synchronised mothers were used to initiate brood size or egg-laying and survival assay, to remove potential effects of maternal age for phenotypic scoring. Highlighted in bold are the starting points for the two assays (2-hour synchronised eggs (P_0_)), or the points of phenotypic readout. Brood size was calculated by summation of the number of F1 adults for the five days.

**Fig. S5:**
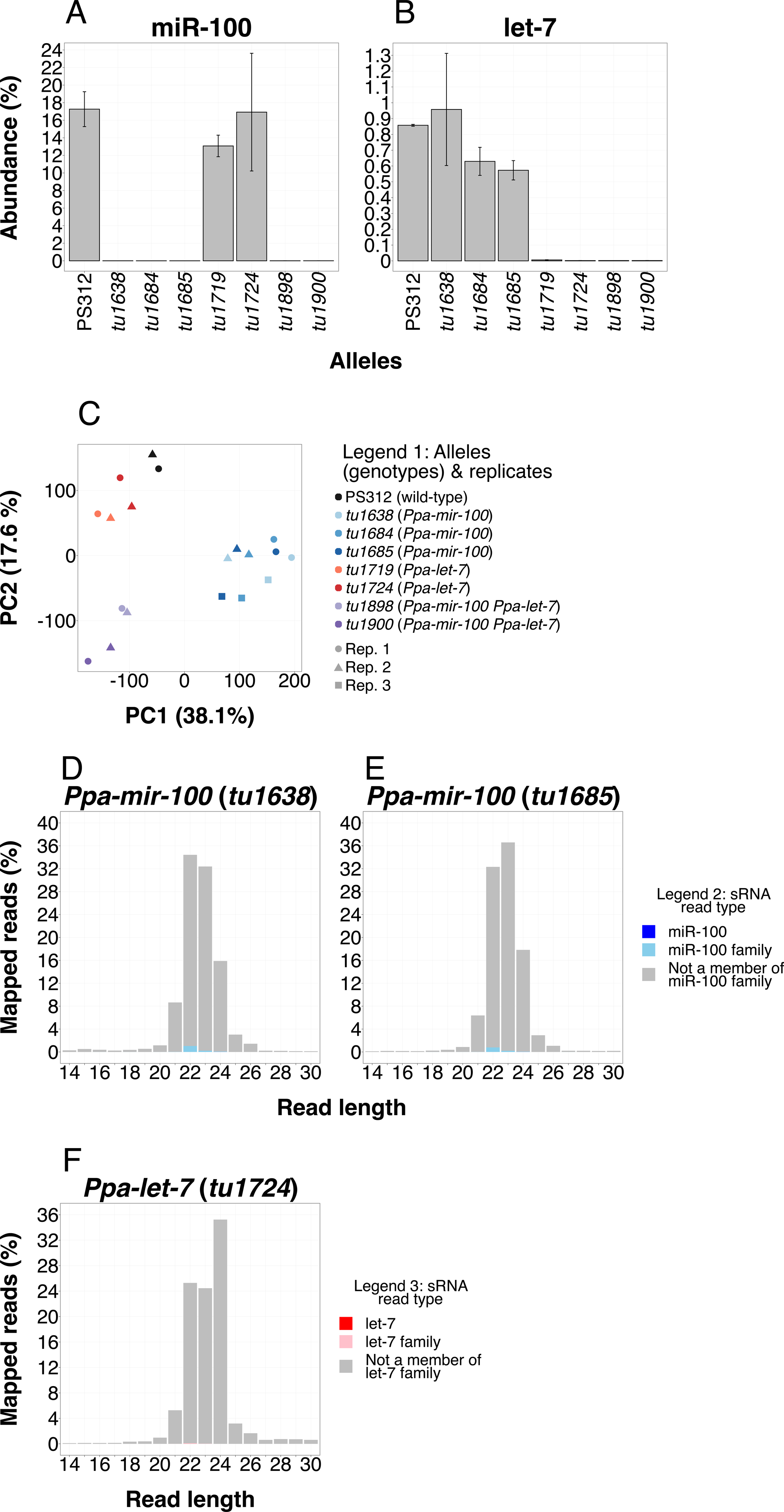
Inter-allelic genotypic congruence among *Ppa-mir-100*, *Ppa-let-7*, and *Ppa-mir-100 Ppa-let-7* double mutants: A–B: Expression levels (mean of three biological replicates for *Ppa-mir-100* alleles, and of two biological replicates for the other three genotypes) of miRNAs miR-100 (A) and let-7 (B) across wild-type and multiple alleles of the three mutant genotypes (*Ppa-mir-100*, *Ppa-let-7*, *Ppa-mir-100 Ppa-let-7*). Error bars represent the SEM. Note that the y-axis is different between panels. C: Principal component analysis (PCA) of the wild-type and mutant alleles of J3 stage-specific small RNA transcriptome. Colours correspond to different alleles, and shapes correspond to different biological replicates per developmental stage (Legend 1). Alleles of the same genotype (mentioned in brackets in Legend 1) are represented with different shades of the same colour (*mir-100*: blue; *let-7*: red; *mir-100 let-7*: purple). D–F: Profile of miR-100 family of miRNAs in *Ppa-mir-100* alleles (D–E) or of let-7 family of miRNAs in *Ppa-let-7*(*tu1724*) (F). Also see Fig. 5F for comparison to panels D and E, and Fig. 5K for comparison to panel F. Legend 2 corresponds to panels D and E, Legend 3 to panel F.

**Fig. S6:**
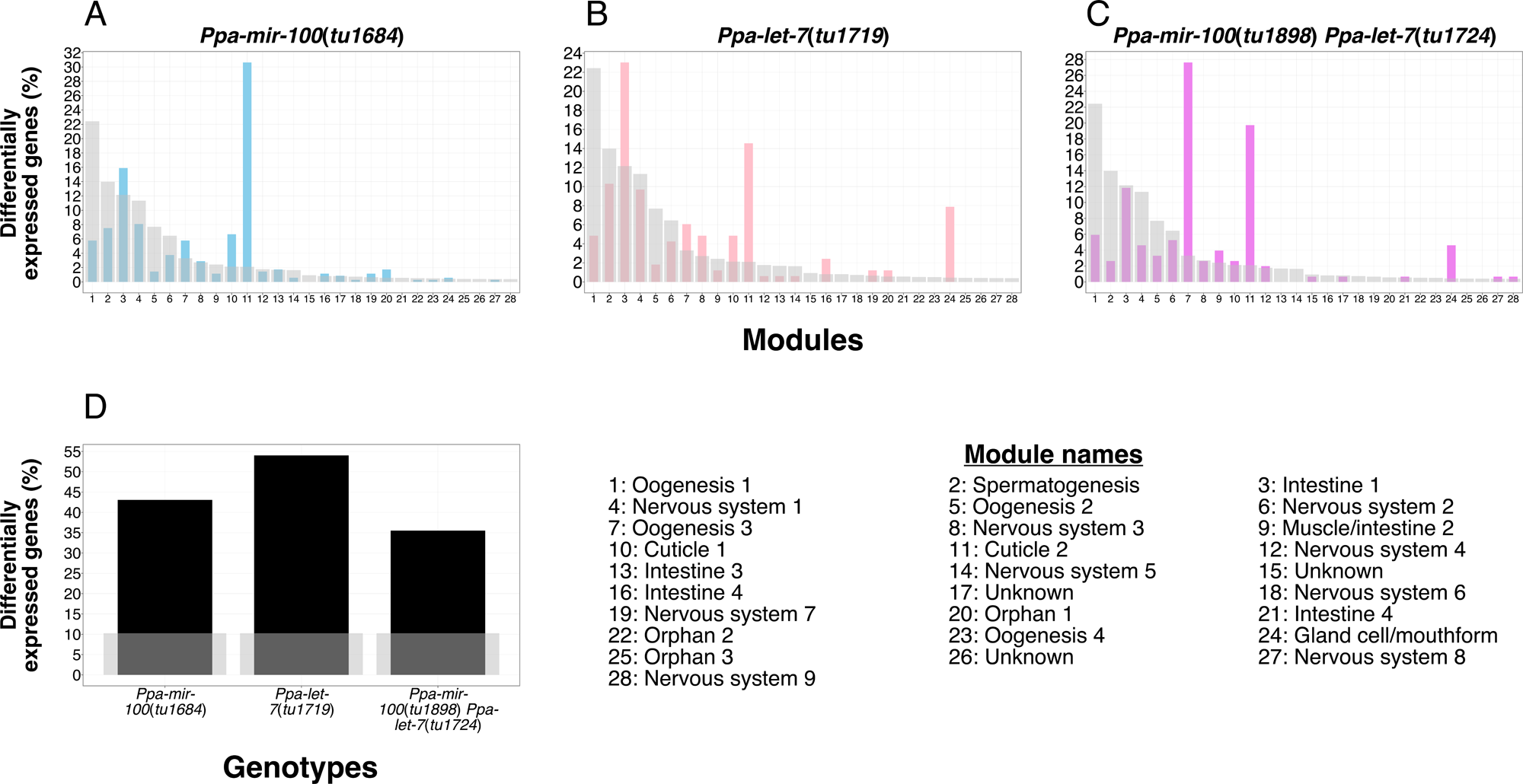
Overrepresentation analyses of *Ppa-mir-100*, *Ppa-let-7*, and *Ppa-mir-100 Ppa-let-7* double mutants of *P. pacificus*: A–C: Gene module overrepresentation analysis of the three mutant genotypes, shown as overlapping bar plots. Translucent grey bars in all three panels represent reference distribution as created in [94], and their corresponding names are also mentioned underneath for reference. Note that the y-axis is different between the three panels. D: Oscillating genes overrepresentation analysis of the three mutant genotypes, shown as overlapping bar plots. Translucent grey bars represent the baseline percentage of oscillating genes in the genome as identified in [93].

**Fig. S7:**
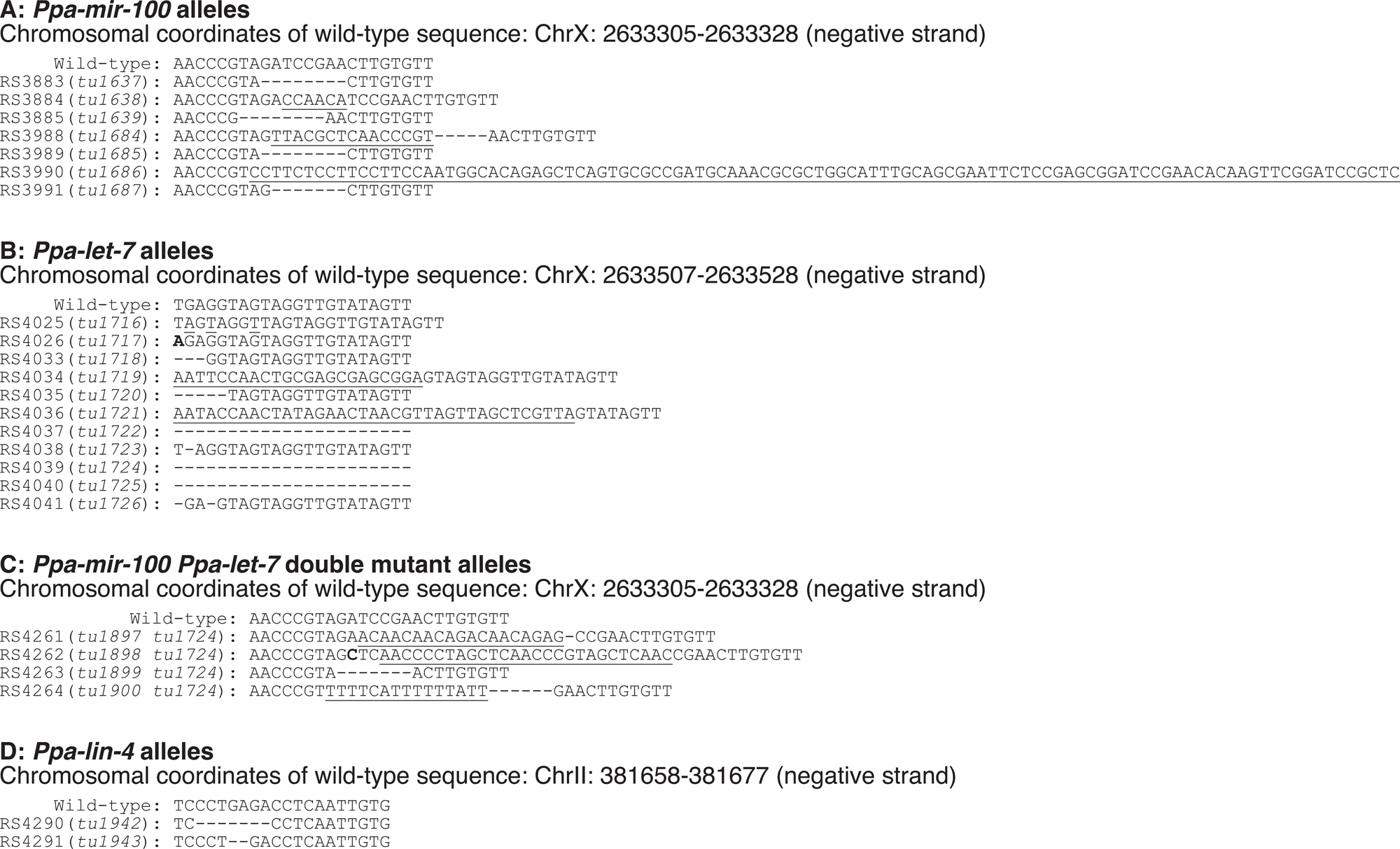
Molecular lesions of all alleles generated in this study (chromosomal coordinates of the corresponding wild-type sequences of miRNAs are according to El Paco V3 assembly of *P. pacificus* genome; deletions are shown by a dash (-), insertions by underlined letters, and SNPs by bold letters): A: *Ppa-mir-100* alleles. Allele *tu1684* has a 15-nucleotide insertion followed by a 5-nucleotide deletion w.r.t. wild-type genomic sequence, and allele *tu1686* has a truncated gene after 7 nucleotides followed by 147-nucleotide insertion w.r.t. wild-type genomic sequence. Note that the alleles *tu1637* and *tu1685* have the same net molecular lesion, despite being acquired from independent experiments. B: *Ppa-let-7* alleles. 5 alleles (*tu1718*, *tu1719*, *tu1720*, *tu1721*, and *tu1726*) have gene truncations at their 5’ end, and 3 alleles (*tu1722*, *tu1724*, and *tu1725*) have complete deletions, *i.e.*, null mutants. C: *Ppa-mir-100 Ppa-let-7* double mutant alleles. All double mutant alleles were generated by knocking out *Ppa-mir-100* in *Ppa-let-7(tu1724)* null mutant background. As such, only molecular lesions pertaining to *Ppa-mir-100* gene are shown. Null mutation of *Ppa-let-7* in all double mutant alleles was confirmed using Sanger sequencing. D: *Ppa-lin-4* alleles

**Fig. S8:**
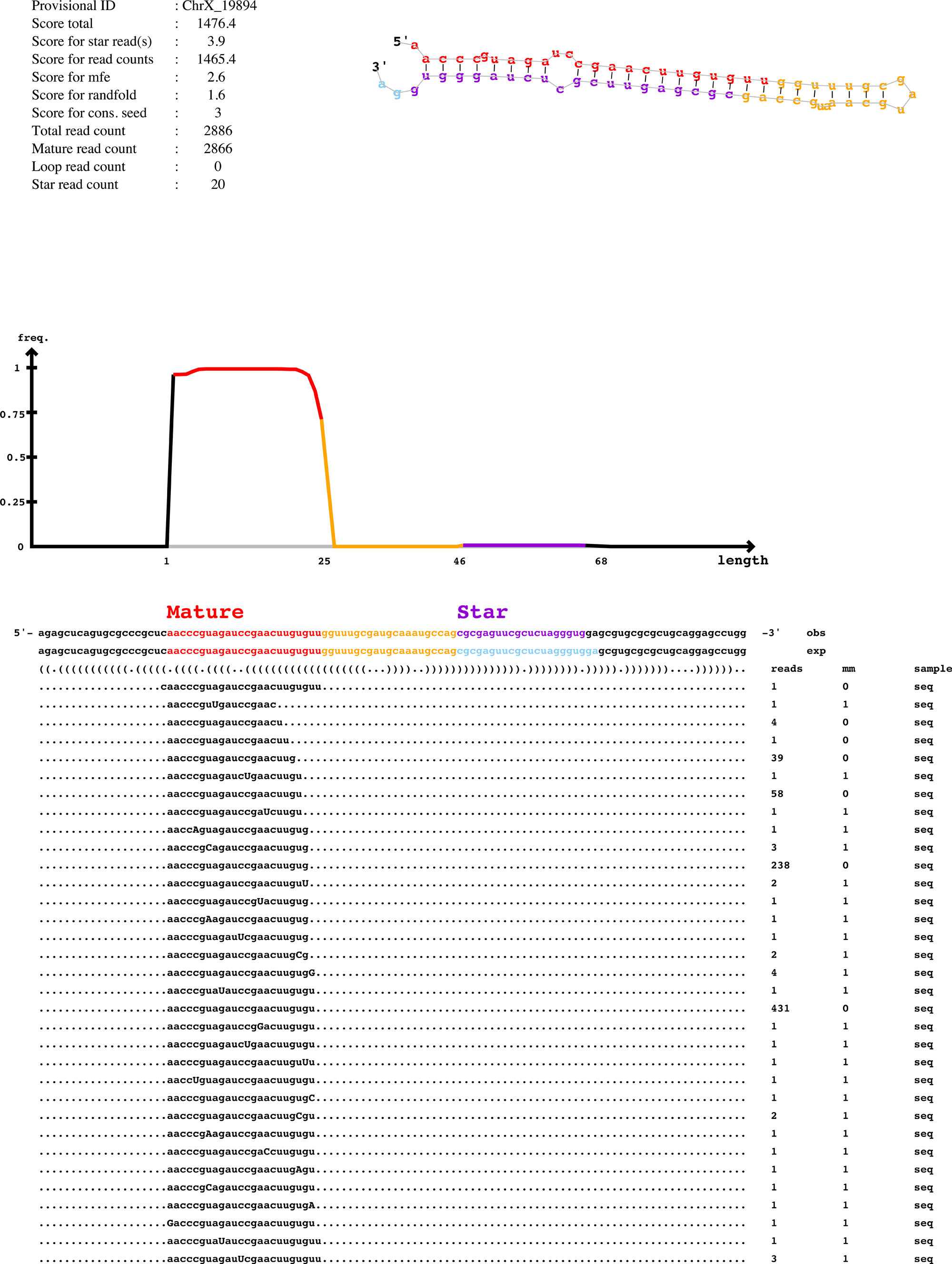
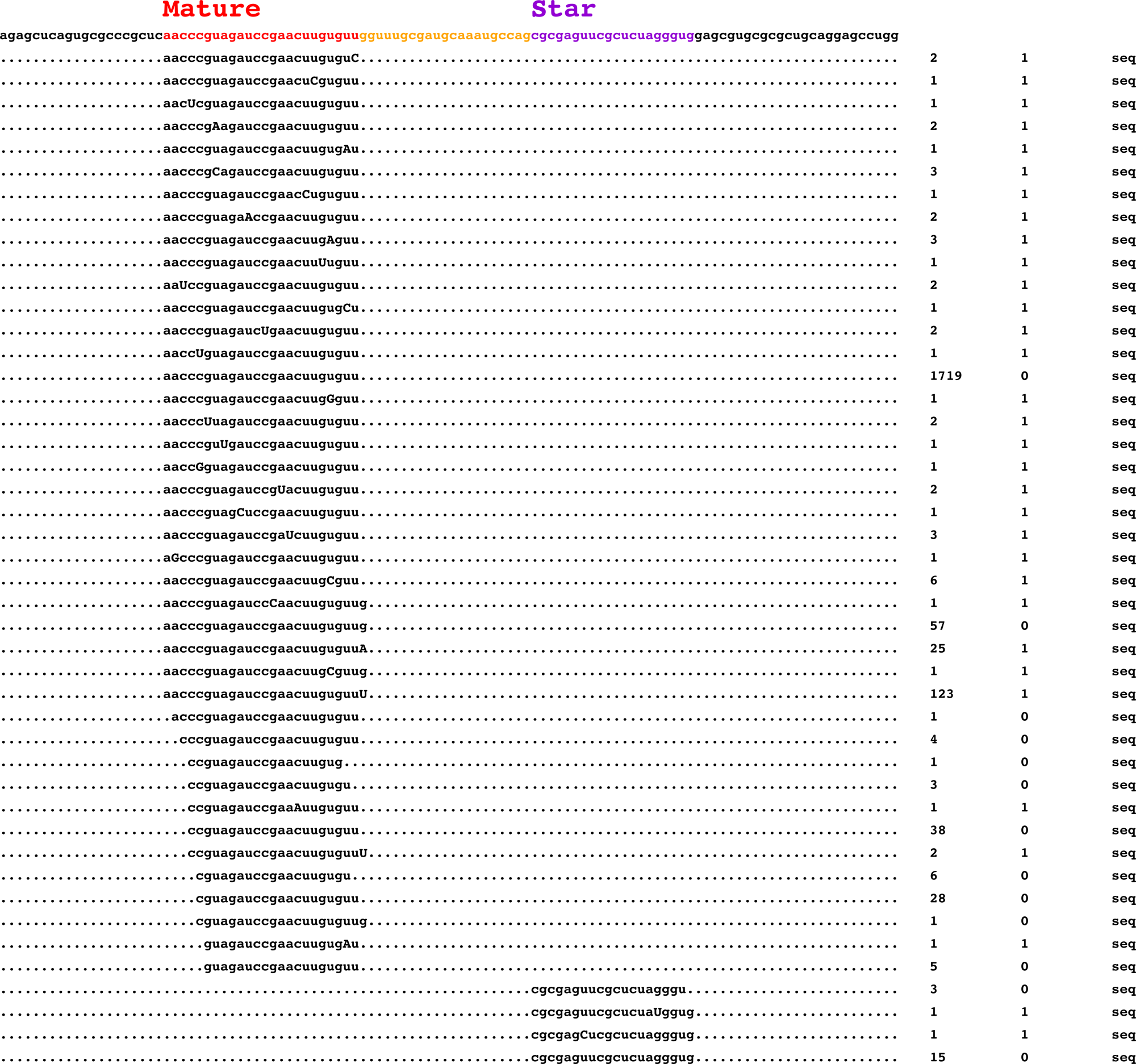
**miR-100 miRDeep2 output** miRDeep2 output corresponding to ppc-miR-100, showing — among other details — the hairpin structure of the corresponding pre-miRNA and 3p miRNA (labelled as ‘Star’ and highlighted in purple)

**Fig. S9:**
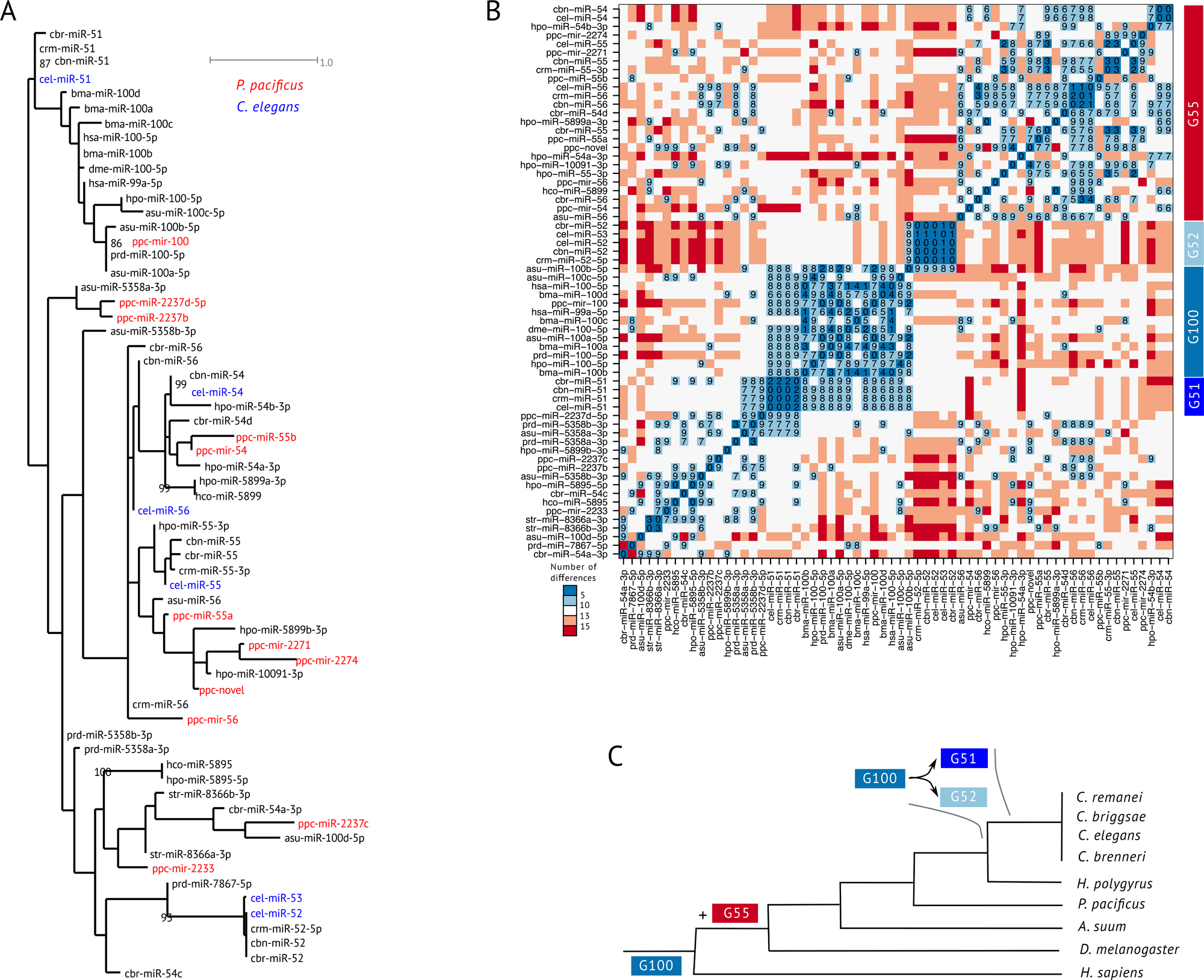
Evolutionary history of the miR-100 family in nematodes: A: We calculated a maximum-likelihood tree of miR-100 family members in nematodes, together with their orthologues in *D. melanogaster* and *H. sapiens*. The tree shows nine members of the miR-100 family in *P. pacificus* (ppc-novel — UACCCGUAGCCCUUCUGCCGCA — was first identified in [41]. Also see Table S5), and suggests that ppc-miR-54, −55, and −56 are orthologues of cel-miR−54, −55, and −56. This would indicate that the expansion of miR-100 family predated the loss of *let-7-complex*. B: Visualisation of the pairwise differences between miR-100 family suggests several orthologous groups — G100, G51, G52, and G55 — within this family. First, all miRNAs annotated as miR-100 by miRBase are less than five differences away from another member of the G100 group. Second, many members of G55 group have at least one other closely related member (number of differences < 5) within the group. Third, groups G51 and G52 form clusters of closely related sequences (number of differences < 5). However, their relationship with regard to other groups is less clear. While G52 members have a weak similarity (number of differences < 10) with a single member of G100, all members of G51 have weak similarity with multiple members of G100, which makes G51 the best candidate for orthology to miR-100 in *Caenorhabditis* nematodes. C: Based on our phylogenetic analysis, we would propose the following scenario for evolution of miR-100 in nematodes: a first expansion likely occurred in the ancestor of analysed nematodes; close to the branch leading to the ancestor of *Caenorhabditis* genus, miR-100 either evolved or was replaced by the groups G51 and G52.

**Fig. S10:**
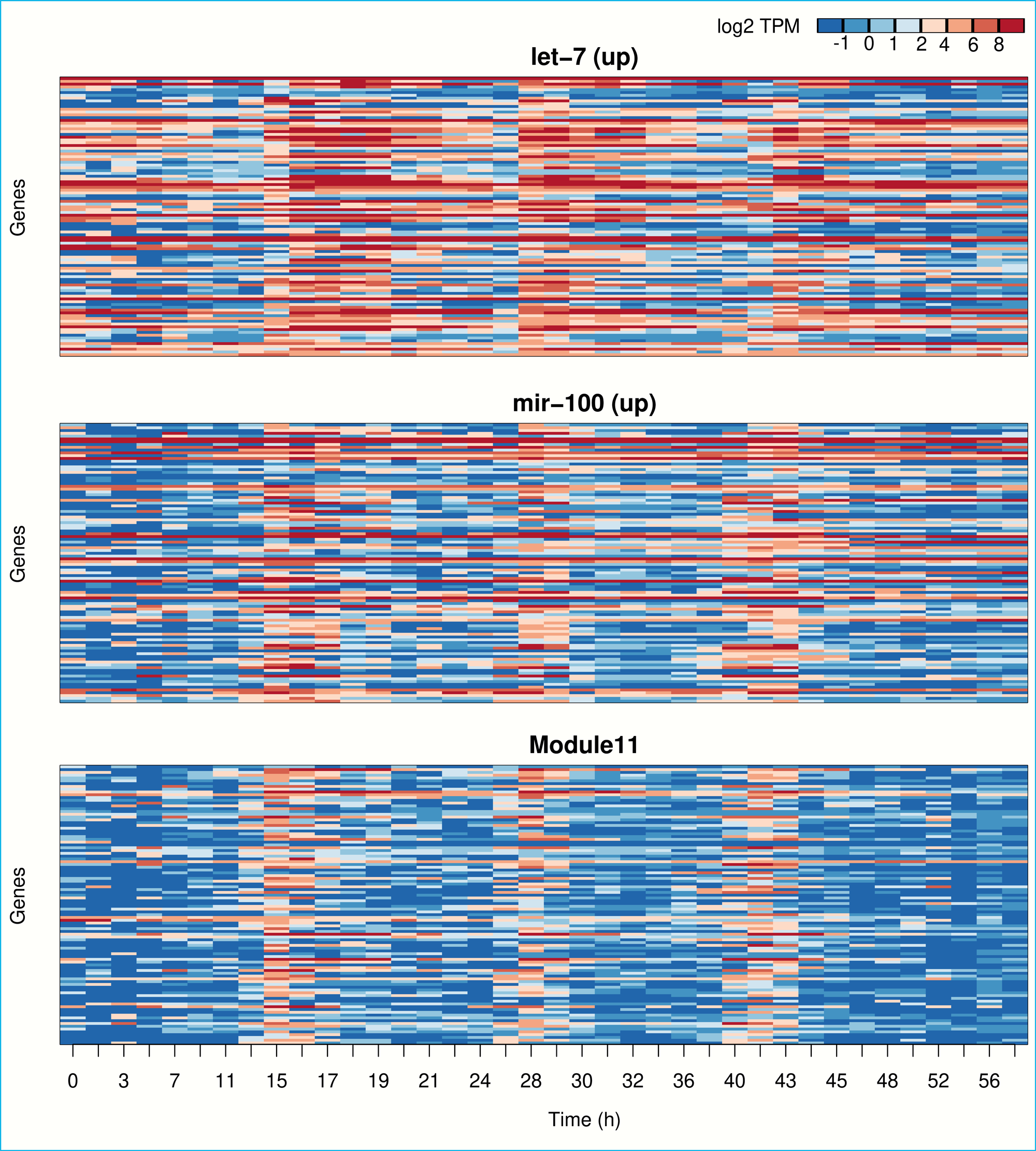
Developmental expression of miRNA-regulated candidate genes: The heat-maps show the expression of 100 randomly subsampled genes from three different gene sets across the development of *P. pacificus* [93]. The first two heat-maps correspond to genes that are upregulated in *Ppa-let-7* and *Ppa-mir-100* respectively. The x-axis shows developmental time post hatching (please refer to [93] for further details). The last panel shows candidate genes of the co-expression module 11 [94] that shows a significant overlap with both sets of differentially expressed genes. All three gene sets show periodic expression patterns with three distinct peaks within the larval development of *P. pacificus*. Although both transcriptomic datasets were sampled at different temporal resolution and are therefore not exactly comparable, the presence of periodic signals in the DEGs suggests that both miRNAs and the genes they regulate downstream are not strictly co-expressed, as the miRNAs are mainly expressed at distinct developmental stages, *i.e.*, miR-100 is the most abundantly expressed at J2 and J3 stages, and let-7 in J3 and J4/YA.

## Declarations

### Ethics approval

Not applicable

### Consent for publication

Not applicable

### Availability of data and materials

All mutant strains generated in this study are available upon request to the corresponding author. The raw sequencing data generated in this study can be accessed at the European Nucleotide Archive (ENA) repository under the accession number PRJEB76785.

## Authors’ contributions

DRS, MSW, and RJS conceptualised the study. DRS and RJS designed the experiments. DRS performed staging and sample collection experiments, RNA extraction and small RNA library preparation for developmental analysis, generated CRISPR mutants (except *lin-4* mutants), performed mutant phenotypic assays, performed all sequencing data analysis except miRDeep2 analysis for identification of novel miRNAs, phylogenetic analysis of miR-100 family members, developmental expression analysis of miRNA-regulated candidate genes, and identification of direct mRNA targets of miRNAs, and created the corresponding visualisations. CR performed miRDeep2 analysis for identification of novel miRNAs, phylogenetic analysis of miR-100 family members, developmental expression analysis of miRNA-regulated candidate genes, and identification of direct mRNA targets of miRNAs, and created the corresponding visualisations. WR performed RNA extraction for mutant analysis. HW generated the *lin-4* CRISPR mutants. Additionally, CR, WR, and HW mentored DRS in sequencing data analysis, wet-lab experiments, and CRISPR experiments respectively. DRS wrote the manuscript, with inputs from CR, MSW, and RJS. RJS acquired funding for the study, provided the resources, and supervised the study.

## Acknowledgements

The authors are grateful to Heike Haussmann for freezing all the mutant strains generated in this study, Heike Budde of the Genome Centre facility for her assistance in performing small RNA sequencing of the developmental stages, and Gabi Eberhardt for her assistance in shipping RNA samples to Novogene Ltd. for sequencing of mutants. Furthermore, the authors thank Tobias Loschko, who, along with WR, mentored DRS in the initial stages of the project in RNA biology and quality control techniques, Metta Riebesell, who, along with HW, mentored DRS in nematode micro-injections, Dr. Wen-Sui Lo and Dr. Marina Athanasouli for their expertise and engaging discussions pertaining to bioinformatic analyses, Dr. Tobias Theska Renahan, Dr. Tess Renahan, and Dr. Adrian Streit for critically reading the manuscript, providing their insights, and for their thoughtful discussions and ideas throughout the course of the study, and members of Sommerlab, in particular Dr. Catia Igreja, for their valuable suggestions.

