## Supplementary tables S8, S9, and S37 for "Developmental small RNA transcriptomics reveals divergent evolution of the conserved microRNA miR-100 and the *let-7-complex* in nematodes"

**A. Statistical significance of difference of brood size among pairwise comparisons**

| Comparison | Statistical significance |
|---|---|
| WT vs *Ppa-mir-100* | ** |
| WT vs *Ppa-let-7* | *** |
| WT vs *Ppa-mir-100 Ppa-let-7* | *** |
| WT vs *Ppa-lin-4* | n.s. |
| *Ppa-mir-100 Ppa-let-7* vs *Ppa-lin-4* | *** |

| Comparison | Statistical significance |
|---|---|
| *Ppa-mir-100* vs *Ppa-let-7* | * |
| *Ppa-mir-100* vs *Ppa-mir-100 Ppa-let-7* | * |
| *Ppa-mir-100* vs *Ppa-lin-4* | ** |
| *Ppa-let-7* vs *Ppa-mir-100 Ppa-let-7* | n.s. |
| *Ppa-let-7* vs *Ppa-lin-4* | *** |

**B. Statistical significance of difference of brood size per day among pairwise comparisons**

| Day | Comparison | Statistical significance |
|---|---|---|
| 1 | WT vs *Ppa-mir-100* | *** |
| | WT vs *Ppa-let-7* | *** |
| | WT vs *Ppa-mir-100 Ppa-let-7* | *** |
| | WT vs *Ppa-lin-4* | *** |
| | *Ppa-mir-100* vs *Ppa-let-7* | n.s. |
| | *Ppa-mir-100* vs *Ppa-mir-100 Ppa-let-7* | n.s. |
| | *Ppa-mir-100* vs *Ppa-lin-4* | * |
| | *Ppa-let-7* vs *Ppa-mir-100 Ppa-let-7* | n.s. |
| | *Ppa-let-7* vs *Ppa-lin-4* | n.s. |
| | *Ppa-mir-100 Ppa-let-7* vs *Ppa-lin-4* | n.s. |

| Day | Comparison | Statistical significance |
|---|---|---|
| 2 | WT vs *Ppa-mir-100* | *** |
| | WT vs *Ppa-let-7* | *** |
| | WT vs *Ppa-mir-100 Ppa-let-7* | *** |
| | WT vs *Ppa-lin-4* | *** |
| | *Ppa-mir-100* vs *Ppa-let-7* | n.s. |
| | *Ppa-mir-100* vs *Ppa-mir-100 Ppa-let-7* | n.s. |
| | *Ppa-mir-100* vs *Ppa-lin-4* | n.s. |
| | *Ppa-let-7* vs *Ppa-mir-100 Ppa-let-7* | n.s. |
| | *Ppa-let-7* vs *Ppa-lin-4* | n.s. |
| | *Ppa-mir-100 Ppa-let-7* vs *Ppa-lin-4* | n.s. |

| Day | Comparison | Statistical significance |
|---|---|---|
| 3 | WT vs *Ppa-mir-100* | n.s. |
| | WT vs *Ppa-let-7* | * |
| | WT vs *Ppa-mir-100 Ppa-let-7* | * |
| | WT vs *Ppa-lin-4* | *** |
| | *Ppa-mir-100* vs *Ppa-let-7* | n.s. |
| | *Ppa-mir-100* vs *Ppa-mir-100 Ppa-let-7* | n.s. |
| | *Ppa-mir-100* vs *Ppa-lin-4* | *** |
| | *Ppa-let-7* vs *Ppa-mir-100 Ppa-let-7* | n.s. |
| | *Ppa-let-7* vs *Ppa-lin-4* | *** |
| | *Ppa-mir-100 Ppa-let-7* vs *Ppa-lin-4* | *** |

| Day | Comparison | Statistical significance |
|---|---|---|
| 4 | WT vs *Ppa-mir-100* | *** |
| | WT vs *Ppa-let-7* | *** |
| | WT vs *Ppa-mir-100 Ppa-let-7* | *** |
| | WT vs *Ppa-lin-4* | *** |
| | *Ppa-mir-100* vs *Ppa-let-7* | n.s. |
| | *Ppa-mir-100* vs *Ppa-mir-100 Ppa-let-7* | n.s. |
| | *Ppa-mir-100* vs *Ppa-lin-4* | *** |
| | *Ppa-let-7* vs *Ppa-mir-100 Ppa-let-7* | n.s. |
| | *Ppa-let-7* vs *Ppa-lin-4* | *** |
| | *Ppa-mir-100 Ppa-let-7* vs *Ppa-lin-4* | *** |

| Day | Comparison | Statistical significance |
|---|---|---|
| 5 | WT vs *Ppa-mir-100* | n.s. |
| | WT vs *Ppa-let-7* | *** |
| | WT vs *Ppa-mir-100 Ppa-let-7* | *** |
| | WT vs *Ppa-lin-4* | *** |
| | *Ppa-mir-100* vs *Ppa-let-7* | * |
| | *Ppa-mir-100* vs *Ppa-mir-100 Ppa-let-7* | n.s. |
| | *Ppa-mir-100* vs *Ppa-lin-4* | *** |
| | *Ppa-let-7* vs *Ppa-mir-100 Ppa-let-7* | n.s. |
| | *Ppa-let-7* vs *Ppa-lin-4* | * |
| | *Ppa-mir-100 Ppa-let-7* vs *Ppa-lin-4* | * |

**C. Statistical significance of difference of frequency of egg-laying among pairwise comparisons**

| Comparison | Statistical significance |
|---|---|
| WT vs *Ppa-mir-100* | *** |
| WT vs *Ppa-let-7* | *** |
| WT vs *Ppa-mir-100 Ppa-let-7* | ** |
| WT vs *Ppa-lin-4* | n.s. |
| *Ppa-mir-100 Ppa-let-7* vs *Ppa-lin-4* | ** |

| Comparison | Statistical significance |
|---|---|
| *Ppa-mir-100* vs *Ppa-let-7* | n.s. |
| *Ppa-mir-100* vs *Ppa-mir-100 Ppa-let-7* | n.s. |
| *Ppa-mir-100* vs *Ppa-lin-4* | ** |
| *Ppa-let-7* vs *Ppa-mir-100 Ppa-let-7* | n.s. |
| *Ppa-let-7* vs *Ppa-lin-4* | *** |

**D. Statistical significance of difference of embryonic lethality among pairwise comparisons**

| Comparison | Statistical significance |
|---|---|
| WT vs *Ppa-mir-100* | *** |
| WT vs *Ppa-let-7* | n.s. |
| WT vs *Ppa-mir-100 Ppa-let-7* | ** |
| WT vs *Ppa-lin-4* | n.s. |
| *Ppa-mir-100 Ppa-let-7* vs *Ppa-lin-4* | *** |

| Comparison | Statistical significance |
|---|---|
| *Ppa-mir-100* vs *Ppa-let-7* | n.s. |
| *Ppa-mir-100* vs *Ppa-mir-100 Ppa-let-7* | n.s. |
| *Ppa-mir-100* vs *Ppa-lin-4* | *** |
| *Ppa-let-7* vs *Ppa-mir-100 Ppa-let-7* | n.s. |
| *Ppa-let-7* vs *Ppa-lin-4* | * |

**E. Statistical significance of difference of survival to adulthood among pairwise comparisons**

| Comparison | Statistical significance |
|---|---|
| WT vs *Ppa-mir-100* | n.s. |
| WT vs *Ppa-let-7* | *** |
| WT vs *Ppa-mir-100 Ppa-let-7* | ** |
| WT vs *Ppa-lin-4* | n.s. |
| *Ppa-mir-100 Ppa-let-7* vs *Ppa-lin-4* | ** |

| Comparison | Statistical significance |
|---|---|
| *Ppa-mir-100* vs *Ppa-let-7* | ** |
| *Ppa-mir-100* vs *Ppa-mir-100 Ppa-let-7* | n.s. |
| *Ppa-mir-100* vs *Ppa-lin-4* | n.s. |
| *Ppa-let-7* vs *Ppa-mir-100 Ppa-let-7* | * |
| *Ppa-let-7* vs *Ppa-lin-4* | *** |

Table S8: Statistical significance of phenotypic differences between genotypes as shown in Fig. 4. Each table corresponds to its namesake figure panel in Fig. 4. Please refer to 'Phenotypic assays' in the Methods section for details.

| El Paco V1 annotation | El Paco V3 annotation | PFAM | Chromosome | $log_2$(FC) *Ppa-mir-100* | $log_2$(FC) *Ppa-let-7* | $log_2$(FC) *Ppa-mir-100 Ppa-let-7* |
|---|---|---|---|---|---|---|
| UMM-S328-9.28-mRNA-1 | PPA05669 | Astacin | X | n.a. | -4.685878953 | n.a. |
| UMM-S2847-7.46-mRNA-1 | PPA42525 | Astacin | IV | n.a. | n.a. | n.a. |
| UMM-S2847-6.45-mRNA-1 | PPA05955 | Astacin | IV | n.a. | -2.375104017 | -1.937456096 |
| UMM-S328-7.47-mRNA-1 | PPA05618 | Astacin | X | n.a. | -3.704532949 | -3.00210115 |
| UMM-S293-8.46-mRNA-1 | PPA16331 | Astacin | X | n.a. | -3.350225515 | -2.867998997 |
| UMM-S328-10.33-mRNA-1 | PPA39735 | CAP | X | n.a. | -1.776499128 | -1.624100142 |
| UMM-S57-4.91-mRNA-1 | PPA32730 | Astacin | I | n.a. | -2.944111154 | -2.263647157 |
| UMM-S328-10.78-mRNA-1 | PPA13058 | CAP | X | n.a. | -1.517549385 | -1.588453213 |
| UMM-S283-11.38-mRNA-1 | PPA39293 | Glyco_hydro_18 | IV | n.a. | -1.6499921 | n.a. |
| UMM-S322-3.5-mRNA-1 | PPA29522 | CAP | X | n.a. | n.a. | n.a. |
| UMM-S293-11.30-mRNA-1 | PPA39470 | CAP | X | n.a. | -2.041405047 | n.a. |
| UMA-S322-3.38-mRNA-1 | PPA21910 | CAP | X | n.a. | -1.475708175 | n.a. |
| UMM-S283-11.45-mRNA-1 | Contig1590-snapTAU.1 | Glyco_hydro_18; MFS_1 | IV | n.a. | -1.686063894 | n.a. |
| UMA-S322-7.39-mRNA-1 | PPA21987 | Astacin | X | n.a. | n.a. | n.a. |
| UMS-S2861-1.50-mRNA-1 | PPA27985 | Astacin | X | n.a. | -2.307702765 | n.a. |
| UMS-S328-0.4-mRNA-1 | PPA30108 | none | X | n.a. | n.a. | n.a. |
| UMS-S10-46.25-mRNA-1 | PPA27560 | none | II | n.a. | -2.378455761 | -2.181107418 |
| UMM-S57-36.5-mRNA-1 | PPA30435 | Lectin_C | I | n.a. | n.a. | n.a. |
| UMA-S2861-1.27-mRNA-1 | PPA34430 | Astacin | X | n.a. | n.a. | n.a. |
| UMM-S250-3.76-mRNA-1 | ppa_stranded_DN18096_c0_g1_i1 | ShK | X | n.a. | n.a. | n.a. |
| UMM-S2857-0.30-mRNA-1 | PPA20266 | Astacin | X | n.a. | -3.158198324 | -2.295116979 |
| UMM-S2857-0.41-mRNA-1 | PPA42924 | Astacin | X | n.a. | -2.810086147 | n.a. |
| UMM-S7-5.16-mRNA-1 | PPA03932 | Astacin | I | n.a. | n.a. | n.a. |
| UMA-S2838-46.74-mRNA-1 | PPA06264 | adh_short; KR; THF_DHG_CYH_C | IV | n.a. | n.a. | n.a. |

Table S9: Overlap between common targets of *Ppa-nhr-1* and *Ppa-nhr-40* as identified in Sieriebriennikov *et. al.*, 2020, and the three miRNA mutant genotypes *Ppa-mir-100*(*tu1684*), *Ppa-let-7*(*tu1719*), and *Ppa-mir-100*(*tu1898*) *Ppa-let-7*(*tu1724*) double mutant

| **Comparison** | **Differentially expressed small RNAs (DEsRNAs)** | | | |
|---|---|---|---|---|
| | **miRNAs** | **piRNAs** | **Others** | **Total** |
| Embryos *vs* J2 | 553 | 17 | 7608 | 8178 |
| Embryos *vs* J3 | 507 | 39 | 16869 | 17415 |
| Embryos *vs* J4/YA | 481 | 248 | 30690 | 31419 |
| J2 *vs* J3 | 84 | 1 | 3128 | 3213 |
| J2 *vs* J4/YA | 507 | 200 | 26685 | 27392 |
| J3 *vs* J4/YA | 111 | 166 | 12359 | 12636 |

Table S37: Summary of differentially expressed small RNAs across the four developmental stages
